# Single nuclei transcriptomics in human and non-human primate striatum implicates neuronal DNA damage and proinflammatory signaling in opioid use disorder

**DOI:** 10.1101/2023.05.17.541145

**Authors:** BaDoi N. Phan, Madelyn H. Ray, Xiangning Xue, Chen Fu, Robert J. Fenster, Stephen J. Kohut, Jack Bergman, Suzanne N. Haber, Kenneth M. McCullough, Madeline K. Fish, Jill R. Glausier, Qiao Su, Allison E. Tipton, David A. Lewis, Zachary Freyberg, George C. Tseng, Shelley J. Russek, Yuriy Alekseyev, Kerry J. Ressler, Marianne L. Seney, Andreas R. Pfenning, Ryan W. Logan

## Abstract

The striatum in the brain is involved in various behavioral functions, including reward, and disease processes, such as opioid use disorder (OUD). Further understanding of the role of striatal subregions in reward behaviors and their potential associations with OUD requires molecular identification of specific striatal cell types in human brain. The human striatum contains subregions based on different anatomical, functional, and physiological properties, with the dorsal striatum further divided into caudate and putamen. Both caudate and putamen are involved in altered reward processing, formation of habits, and development of negative affect states associated with OUD. Using single nuclei RNA-sequencing of human postmortem caudate and putamen, we identified canonical neuronal cell types in striatum (*e.g.,* dopamine receptor 1 or 2 expressing neurons, D1 or D2) and less abundant subpopulations, including D1/D2-hybrid neurons and multiple classes of interneurons. By comparing unaffected subjects to subjects with OUD, we found neuronal-specific differences in pathways related to neurodegeneration, interferon response, and DNA damage. DNA damage markers were also elevated in striatal neurons of rhesus macaques following chronic opioid administration. We also identified sex-dependent differences in the expression of stress-induced transcripts among astrocytes and oligodendrocytes from female subjects with OUD. Thus, we describe striatal cell types and leverage these data to gain insights into molecular alterations in human striatum associated with opioid addiction.

## Main

Rates of deaths from opioid overdose and people diagnosed with opioid use disorder (OUD) are continuing to rise in the United States^1^. Efforts to develop new treatment strategies and to bolster avenues of existing treatments for opioid addiction require a deeper understanding of the changes that occur in the human brain with chronic opioid use. To date, few studies have investigated the cellular and molecular changes in the human brain associated with OUD^2–7^. Recently, we reported alterations in neuroinflammation, among other pathways, in postmortem brain regions from subjects with OUD^2, 5^. Consistent with changes in inflammatory signaling, markers for microglia were also significantly enriched in the striatum of subjects with OUD.

Other pathways related to inflammatory states have been associated with OUD, including dysfunctional aggregation of the tau protein^7, 8^, augmented markers of DNA damage^9^, and increased oxidative stress^10, 11^. Activation of inflammatory signaling is initiated by a myriad of processes in the brain. For example, elevated DNA damage can lead to the activation of microglia to release proinflammatory cytokines^12, 13^. Proinflammatory states in the brain can lead to accumulation of reactive oxygen species and metabolic alterations^14–1612,13^in glia and neurons^13, 17–19^. DNA damage, accumulation of tau protein, and neuroinflammation are also implicated in early pathogenesis of neurodegenerative disorders^0–2317, 24, 25^. Thus, accumulating evidence from the human brain suggests alterations in pathways involved in inflammation, oxidative stress, and DNA damage, including several factors associated with neurodegenerative disorders. Further investigating these molecular alterations associated with OUD using single cell technologies may yield important insights into the potential roles for DNA damage and neuroinflammation in specific cell types in the human brain.

Single cell RNA-based approaches in the human brain provide high-throughput analyses to characterize the transcriptomes of specific neural cell types. Studies using single nuclei RNA-sequencing (snRNA-seq) may further resolve disease-associated molecular alterations to specific neural cell types previously found using other approaches (e.g., bulk tissue RNA-seq). By stratifying pathways across different cell types, an aim is to also unmask new molecular pathways providing deeper insights into mechanisms specifically in glial and neuronal cell types associated with disease. A brain region heavily implicated in OUD, along with other psychiatric and neurological disorders, is the striatum. In the human brain, the striatum is anatomically divided into the dorsal and ventral striatum, further subdivided into the nucleus accumbens, caudate and putamen. Functional changes in the caudate and putamen of the dorsal striatum are involved in altered reward processing, habitual drug-seeking and relapse, along with the development of negative affect during opioid withdrawal^26, 29, 2728^, underlying the key roles of the dorsal striatum in OUD. However, only a few studies have examined specific cell types in human dorsal striatum^30^, and to date, no study has resolved the cell type-specific transcriptional alterations in the human striatum associated with OUD.

To begin to investigate OUD-associated alterations across striatal cell types, we conducted snRNA-seq on the dorsal striatum from human postmortem brain and compared the cell type-specific molecular signatures between unaffected subjects and subjects with OUD (98,848 total nuclei from the caudate and putamen of males and females, 12 subjects and 24 biological samples). Given the relatively high number of nuclei per sample, we were able to identify heterogeneous striatal cell types based on anatomical and transcriptional profiles. By comparing unaffected and OUD subjects, we found significant molecular alterations in neuroinflammatory, metabolic, and DNA damage-related pathways, further implicating neuronal generated DNA damage response and microglial activation associated with OUD in the human striatum^7, 31–36^. In postmortem human brains, we observed an elevated enrichment of DNA damage markers, specific to neuronal subtypes. To further assess the relationships between OUD and DNA damage markers, we examined the expression of DNA damage markers across striatal cell types in rhesus macaques that had been administered opioids chronically (∼6 months of twice daily administration). Notably, elevated DNA damage markers identified in OUD subjects were also found in specific striatal neuronal cell types in the rhesus macaque brain. Additionally, we explored the interaction between diagnosis and sex, finding sex- and cell type-specific molecular changes associated with OUD. As a resource, we deposited the annotated and filtered single nuclei transcriptomes for human caudate and putamen in unaffected and OUD subjects on the <u>CZ CELLxGENE Discover Portal</u>.

## Results

### High quality single nuclei transcriptomics identifies canonical and low abundant cell types in human dorsal striatum

To investigate the molecular features of human dorsal striatum, we collected both caudate and putamen from unaffected subjects (n=6, 3 females and 3 males, for each subregion). Unaffected subjects were then compared to subjects with OUD to identify cell type-specific molecular alterations associated with opioid addiction. Both striatal regions were dissected from frozen postmortem tissue samples and processed to generate nuclei suspensions for snRNA-seq (Figure 1A,B). Quality control analysis yielded a total of 98,848 nuclei for subsequent analyses–∼70% of overall nuclei captured were used for analysis with an average of 5,757 and 3,440 nuclei per caudate and putamen, respectively, yielding an average of 8,237 nuclei per subject (Figure 1C). Across subjects, the number of nuclei captured and percent of nuclei analyzed were consistent (p=0.40, linear regression, Table S1). Each nuclei was deeply sequenced at 70% saturation rates, detecting an average of 3,513 ± 77 genes (mean ± standard error) and 13,635 ± 560 unique transcripts per nucleus per subject (Figure 1C). The average number of genes detected and the number of unique mapped identifiers for transcripts were consistent between unaffected and OUD subjects (p=0.55, linear regression, Table S1).

**Figure 1.**
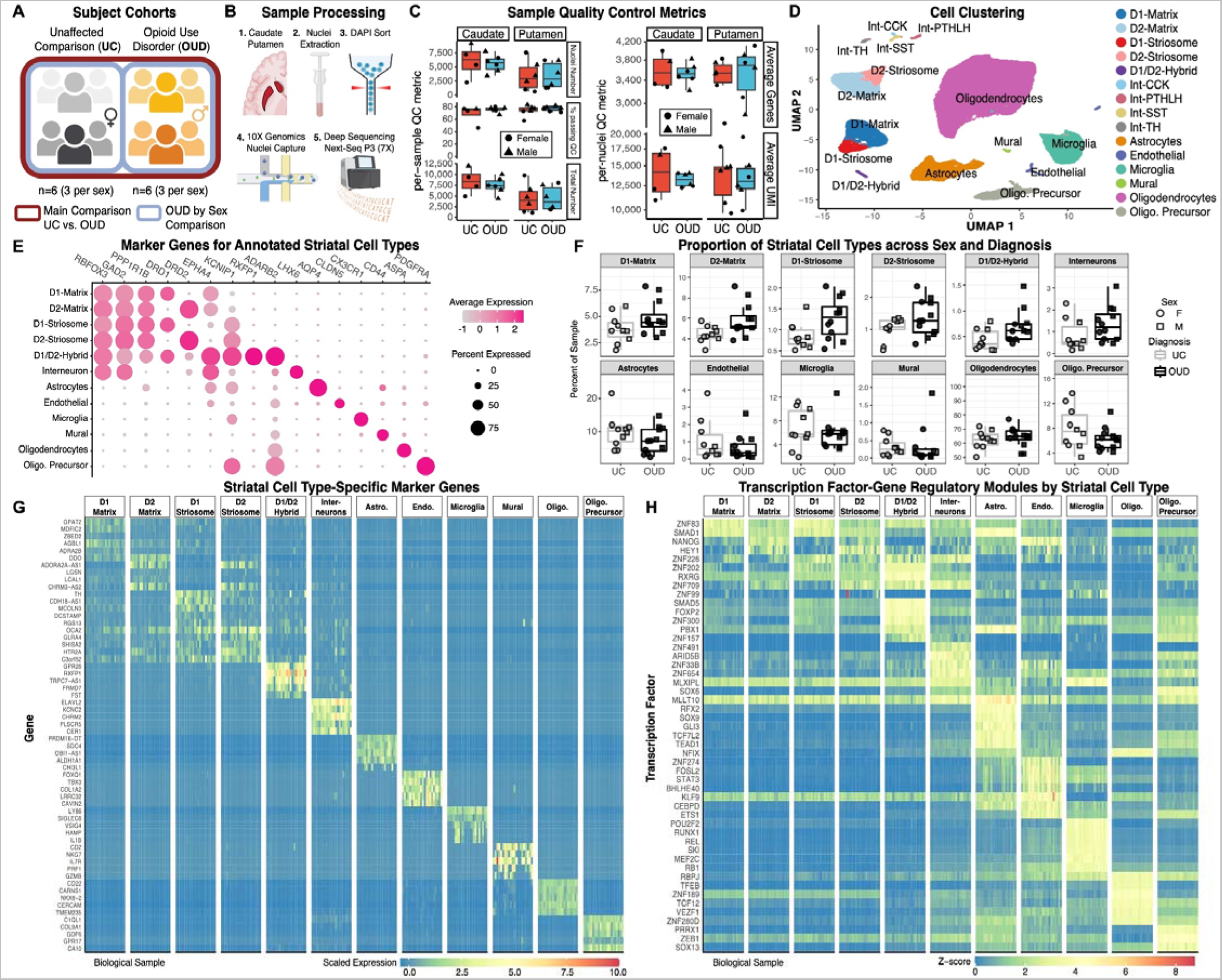
Single nuclei RNA-sequencing of postmortem brain to identify specific cell types in human striatum. (A) Post-mortem human brain cohort design and analysis to compare unaffected comparison (UC) subjects with subjects with opioid use disorder (OUD). (B) Schematic of the sample collection process to isolate single nuclei and generate single nuclei RNA-seq libraries in balanced batches from the caudate nucleus and putamen. (C) Per-sample quality control (QC) metrics and (E) average per-nuclei QC metrics across tissue sources and diagnoses. (D) Single nuclei clustering and label annotation of dorsal striatal cell types using a high quality reference^37^. A low dimensionality projection of striatal cell types after QC filtering and annotation. (E) Dot plots of the marker genes used to annotate the various cell types of the dorsal striatum based on a non-human primate reference. The normalized expression patterns are averaged across all cells and individuals. (F) Boxplot showing the relative percent of each cell type detected in each biospecimen that passes quality control. No significant differences were observed between unaffected comparison and OUD individuals. (G) Cell type-specific marker genes and (H) marker transcription factor-gene regulatory networks discovered from single nuclei RNA-seq striatal tissues.

We used multiple rounds of graph-based clustering, cell label transfers and refinement from a high-quality, high-resolution snRNA-seq non-human primate striatum dataset^37, 38^ (Figure 1D, S1, Methods). The first round identified major cell classes of glia and neuron (primary marker): astrocytes (*AQP4*+), endothelial cells (*CLDN*+), microglia (*CX3CR1*+), mural cells (*CD44*+), oligodendrocytes (*ASPA*+), oligodendrocyte precursors (OPC; *PDGFRA*+), and neurons (*RBFOX3*+, *GAD2*+) (Figure 1D,E). The second round identified major striatal neuronal subtypes, including medium spiny neurons (MSN; *PPP1R1B*+) and interneurons (*LHX6*+) (Figure 1D,E). Among MSNs, we identified various canonical subtypes, consistent with our non-human primate dataset^37^, including *DRD1*+ and *DRD2*+ neurons accompanied by markers that distinguish neurochemical compartments, the striosome (*KCNIP1*+) and matrix (*EPHA4*+). We also identified less abundant neuronal subtypes including MSNs expressing both *DRD1* and *DRD2* (D1/D2-hybrid, known as eccentric MSNs, D1-*Pcdh8^+^*, or D1H in mice)^19, 37–40^, mural cells, and several types of interneurons (*e.g.*, *CCK*+, *PVALB*+, *SST*+, and *TH*+; (Figure 1D,E)^41, 42^. D1/D2-hybrid MSNs expressed conserved marker genes largely distinct from D1- and D2-MSNs (Figure 1E, S2S). These D1/D2-hybrid MSNs co-express both *DRD1* and *DRD2* transcripts at the single cell level (Figure S3), consistent with *in situ* mRNA hybridization and single cell RNA-seq data from both mouse^38, 39, 43^ and non-human primates^37^. The average per cell type metrics showed more genes and unique transcripts in neurons (Number of genes_Neurons_ = 7,318 ± 125; UMsI_Neurons_ = 44,585 ± 1,301), compared to glial cells (Number of genes_Neurons_ = 2,911 ± 74; UMsI_Neurons_ = 8,433 ± 349; Figure S4). Further, proportions of low or high abundance striatal cell types were consistent across brain regions, between sexes, and between unaffected and OUD subjects (FDR=0.25, mixed effects linear regression; Figure 1F; Table S2). Overall, we captured a high-quality snRNA-seq dataset of the postmortem human caudate and putamen that is deeply sequenced and annotated for the study of opioid use disorder.

### Neuronal and glial cell type-specific expression of opioid receptors in human striatum

Opioids, including endogenous opioids, enkephalin, dynorphin, and β-endorphin, differentially activate several classes of opioid receptors, including mu, delta, and kappa, encoded by *OPRM1, OPRD1,* and *OPRK1*, respectively. Previous efforts have mapped striatal expression of opioid receptors in non-human primates and rodents, characterizing species-specific and cell type-specific patterns of expression^44^. We investigated the expression of opioid receptors and cognate ligands within the dorsal striatum.. First, *OPRM1* was detected in each MSN subtype, with the highest level of expression in D1/D2-hybrid and D2-striosome MSNs (Figure S5). We reproducibly detected *OPRM1* in microglia across subjects and biological samples, consistent with findings in rodents^35, 45^, establishing a potential pathway for opioids to directly modulate neuroimmune processes^5^. D1-striosome MSNs expressed the gene encoding the preproprotein, prodynorphin (*PDYN*), while D2-matrix and D2-striosome MSNs coexpressed *OPRD1* and proenkephalin (*PENK*) at markedly higher levels than other striatal cell types (Figure S5). Preferential expression of *OPRD1* and *PENK* in D2 MSNs in human striatum is consistent with rodent striatum^46^. Across MSNs, we detected *OPRK1* expression at many folds lower than *OPRM1*, *OPRD1*, *PENK*, or *PDYN*, suggesting that targets of the dynorphin-kappa opioid receptor signaling by D1-striosome MSNs may primarily be outside of dorsal striatum^47, 48^. Between unaffected and OUD subjects, we found no significant differences in opioid receptor and/or endogenous ligand expression within specific cell types. Other mechanisms involved in the post-transcriptional and/or post-translational regulation of opioid receptor and ligand expression, along with receptor binding activity, may be altered in OUD.

### Identification of cell type-specific transcription factor and gene regulatory networks in human dorsal striatum

Following cell annotation, we aimed to identify putative regulatory transcription factor-gene regulatory networks across striatal cell types. First, we identified marker genes for each of the annotated cell types that were reproducible across biological replicates (mean ± standard error, 722 ± 172 specific marker genes per cell type; Figure1D,E,G, S1; Table S3). Glial cell types yielded more marker genes (1,134 ± 204) compared to neuronal cell types (310 ± 131). Next, we inferred gene regulatory networks directly from each of the cell type-specific marker datasets using machine-learning (SCENIC)^49^. SCENIC is based on the premise that transcriptional regulation of gene expression is, in part, based on transcription factor binding to proximal gene promoters. Based on single cell gene expression and the enrichment of promoter binding sites, we used SCENIC to build transcription factor gene regulatory modules that were specific to each of the neural cell types identified in human striatum. Collectively, we found many transcription factors and gene regulatory modules per cell type (47 ± 15 specific modules; glial subtypes, 90±19; neuronal subtypes, 12 ± 6; Figure 1H; Table S4). For example, ZNF83 (HPF1) transcription factor-regulatory module was enriched in D1-matrix and D1-striosome MSNs, known to be involved in DNA damage repair. RXRG^30, 50, 51^ and FOXP2^52^ transcription factor-regulatory modules were highly enriched in D1/D2-hybrid MSNs relative to other MSN subtypes, both of which are implicated in psychiatric disorders, including substance use^53^. ZNF202 was also highly enriched in D1/D2-hybrid MSNs–transcription factor involved in cellular metabolism and the direct modulation of dopamine receptor 3 (DRD3). Other modules included SOX9^54^ and TCF7L2^55^ enriched in astrocytes, and others in microglia, such as RUNX1, REL, and MEF2C, with the RB1 and NFIX^56^ modules enriched in oligodendrocytes (Figure 1H). Together, the annotated cell type clusters in human dorsal striatum, their associated marker genes, and related transcription factor gene regulatory modules offer new insights into the molecular signaling pathways that contribute to the heterogeneity of cellular identity in human striatum.

### Differentially expressed genes in specific striatal cell types associated with OUD

We assembled a cohort of subjects diagnosed with OUD based on time since clinical diagnosis of at least four years and deceased from opioid overdose, similar to our prior work^4, 5^. OUD subjects were matched with unaffected subjects on sex, age, postmortem interval (PMI), and RNA integrity number (RIN) (Table S5). Gene expression profiles were aggregated across cell types to generate pseudobulk profiles for each major cell type. To identify differentially expressed genes (DEGs), gene expression profiles were compared between unaffected and OUD subjects across cell types, while also considering covariates (age, PMI, RIN, cell type abundance, gene detection rate, and surrogate variables). We identified 1,765 DEGs across cell types (197 ± 43 genes per cell type; FDR < 0.05; Figure 2; Table S6), with more DEGs found in glial cells relative to neurons (Figure 2A,B).

**Figure 2.**
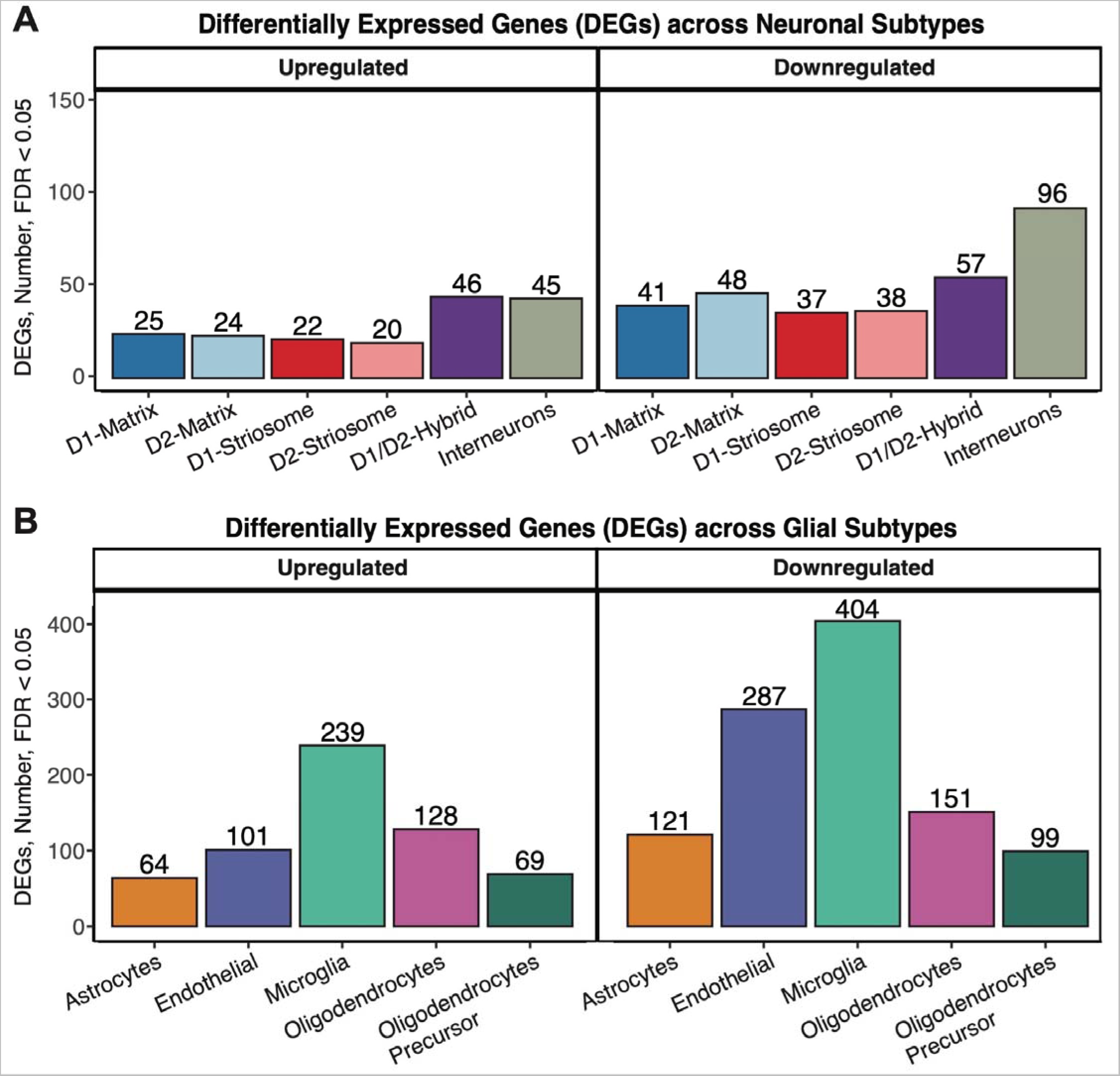
Differentially expressed genes in specific neuronal and glial cell types in human striatum associated with OUD. (A) Barplot of differentially expressed genes (DEGs) either up- or down-regulated in striatal neurons at a false discovery rate less than 0.05 (FDR < 0.05). (B) Barplot of differentially expressed genes (DEGs) either up- or down-regulated in striatal glia at a false discovery rate less than 0.05 (FDR < 0.05).

Using gene set enrichment analyses (GSEA), we identified pathways significantly enriched in aggregate cell types and across specific cell types. Across cell types, we found 286 pathways (34 ± 7 pathways per cell type; FDR < 0.05, Table S7) and 51 differentially regulated transcription factor regulatory modules (5.1 ± 0.7 per cell type, FDR < 0.05, Table S8). The cell type-specific transcriptional differences in OUD subjects were represented largely by distinct biological processes between neurons and glia (Figure S6).. Among MSNs, DEGs had a tendency to be shared between MSNs, while DEGs tended to be more distinct between glial subtypes and between interneuron subtypes (Figure S5). We identified a higher number of DEGs in interneuron subtypes and D1/D2-hybrid MSNs compared to canonical D1- or D2-MSNs. DEGs in D1/D2-hybrid MSNs were completely unique relative to canonical MSNs, suggesting subpopulations of striatal MSNs exhibit different transcriptional responses to chronic opioid use and factors associated with OUD (Figure S5).

### Enriched pathways among neurons converge on various pathways of cell stress in OUD

We conducted separate pathway analyses on upregulated and downregulated transcripts (Figure 3A) between unaffected and OUD subjects. In neurons, most of the upregulated pathways in OUD were related to processes of cell stress response (Figure 3C). For example, pathways of DNA replication and cell cycle re-entry were upregulated in MSNs (Figure 3C; Table S9)^57^. Both *MCM8* (log2FC > 1.06, FDR < 0.041) and myosin light-chain 6, *MYL6*, (log2FC >1.13, FDR < 0.045), genes involved in DNA replication, were significantly upregulated in striatal MSNs (Figure 3A). MCM8, forms a complex with MCM9 to facilitate repair of double-stranded DNA breaks^58, 59^. MYL6 is a key factor in rho-GTPase signaling, known to facilitate glutamate receptor endocytosis^60^ and neuroplasticity^61, 62^, and may be involved in DNA damage via the activation of RAC1^63^. *APOE*^64–66^ (striosome MSNs and D2-matrix MSNs: log2FC >2.50, FDR < 0.025; Figure 3A) and *GPX4* (D1-striosome MSNs: log2FC>1.17, FDR < 0.044) were also upregulated in MSNs of OUD subjects. Both *APOE* and *GPX4* are involved in neuronal oxidative stress, where GPX4 buffers the accumulation of reactive oxygen species to prevent apoptosis^64–66^. Accumulation of reactive oxygen species and oxidative stress in neurons involves alterations in redox signaling and mitochondrial respiration^67^. Indeed, across multiple MSN subtypes, genes involved in mitochondrial respiration were significantly upregulated in OUD, including a complex III subunit of the mitochondrial respiratory chain, *UQCR11* (log2FC >1.36, FDR < 0.042), and a subunit of complex I, *NDUFA4* (log2FC>1.29, FDR < 0.030).

**Figure 3.**
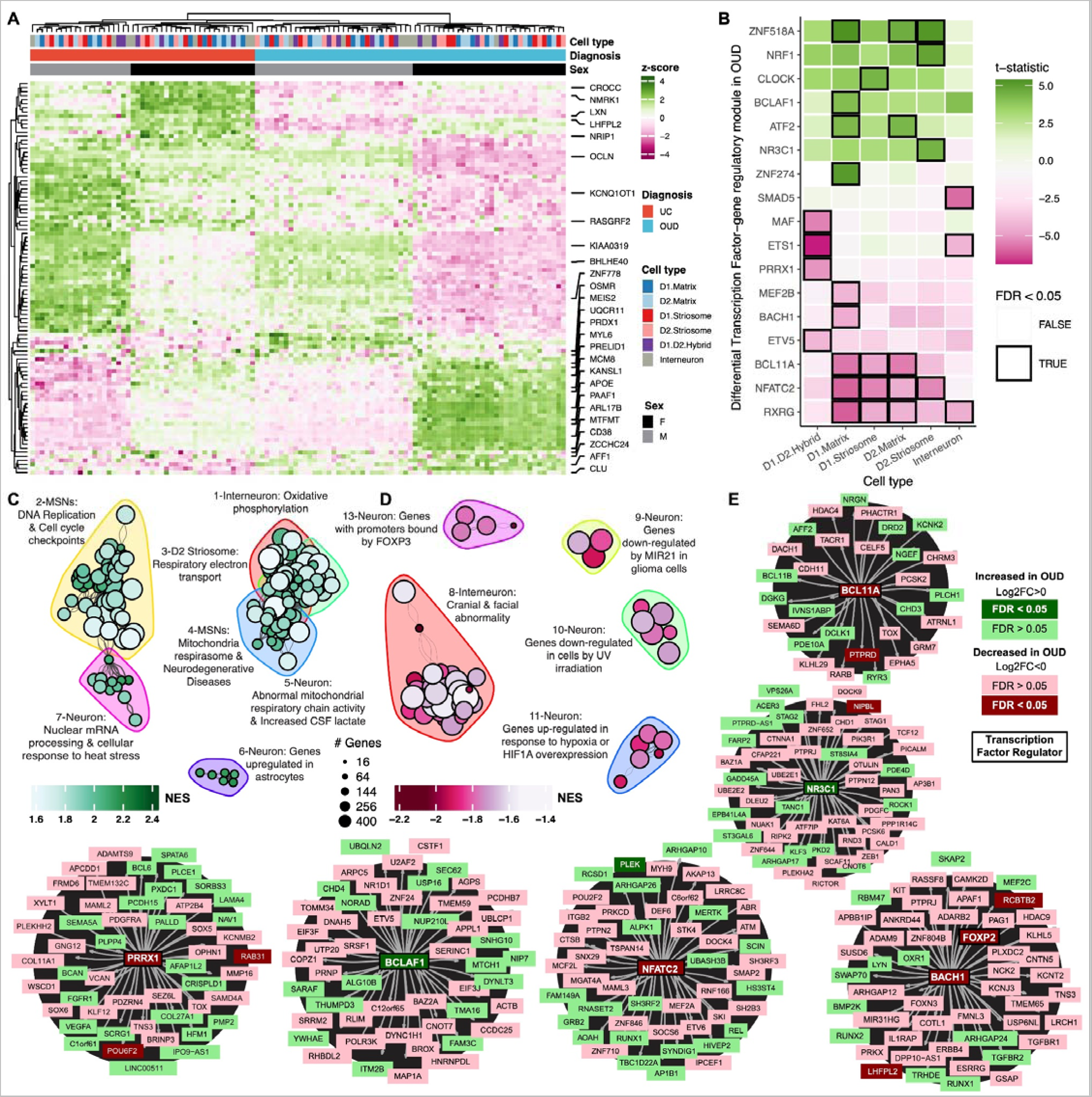
Transcriptional alterations in specific neuronal cell types in human striatum associated with OUD. (A) Expression heatmap of z-normalized pseudobulk gene expression, where the rows are DEGs and the columns are the biological replicates labeled by cell type, diagnosis, and sex of each subject. Genes that are represented by the enriched pathways in (C-D) are labeled along the right margins. (B) A heatmap plotting the t-statistic of differentially activated transcription factor gene regulatory networks across neuronal subtypes in OUD. (C-D) Network plot of significant clusters and enriched pathways, where each point is a significantly enriched pathway in a neuronal cell type and lines represent the proportion of shared genes between two pathways. Each point is colored by the normalized enrichment score (NES), indicating whether a pathway is enriched in up- or down-regulated genes. The color outlines represent the unique clusters of interconnected pathways by shared genes and the nearby text labels the unique cluster number and briefly summarizes which cell types and pathways are represented. (E) Cluster maps of select transcription factor gene regulatory networks. Each transcription factor or gene is colored to denote the direction and significance of DE in neuronal subtypes in OUD.

We identified several transcription factor gene regulatory modules that were significantly upregulated in different neuronal subtypes linked to cell senescence, DNA damage, and inflammation, and stress (Figure 3B,E; FDR < 0.05). For example, the module linked to BCLAF1 was upregulated in OUD, a transcription factor involved in the transduction of NFkB-dependent signaling and the activation of DNA-damage-induced senescence^69, 4268^ (Figure 3E). Another module was associated with the upregulation of the neuron-specific glucocorticoid receptor transcription factor, NR3C1, previously linked to the impact of psychosocial stress on the human brain^70–72^ (Figure 3E). Other stress-related transcription factor gene regulatory modules were upregulated in OUD, including ATF2^73^ and ZNF518A^74^. Both transcription factor modules were upregulated primarily in D1-matrix and D2-matrix MSNs (FDR < 0.04). ATF2 is induced by stress in mouse dorsal striatum^73^ and phosphorylated downstream of delta opioid receptor activation^75^, suggesting delta opioid receptor activation of ATF2-dependent transcription in striatal matrix MSNs^76, 77^ may be due to an interplay between stress and opioids in OUD.

Compared to upregulated modules, we identified almost twice as many significantly downregulated transcription factor gene regulatory modules in OUD subjects (Figure 3D). We identified several downregulated pathways involved in neuroprotection (Figure 3D). For example, the humanin-like 8 gene, *MTRNR2L8*, is significantly downregulated in OUD across every neuronal subtype (log2FC <-3.79, FDR < 0.0054). *MTRHR2L8* and the mitochondrial paralog, *MTRNR2*, act as neuroprotective factors in response to neurodegeneration and the induction of DNA damage^24, 78, 79^. The top DEGs within clusters of pathways involved in UV irradiation, another potential link to DNA damage (FDR < 0.029, Figure 3D), included *FOXO1*, *TLE4*, *SHOC2*, *MEIS2*, and *THBS1* (log2FC <-0.64, FDR < 0.0422) in interneurons and CLASP1, PIK3C3, and MTUS1 (log2FC <-0.64, FDR < 0.042) in D1/D2-hybrid MSNs. Additionally, downregulation of MEF2B and BACH1 transcription factor gene regulatory modules were found in D1-matrix MSNs in OUD (Figure 3E), accompanied by downregulation of MAF, EST1, ETV1, and PRRX1 modules in D1/D2 hybrid MSNs (FDR < 0.043), each of which are implicated in neuronal stress^80–83^. Notably, BACH1, BCL11A, and PRRX1 have downstream targets with concordant differential expression in OUD (*e.g., FOXP2*, *POUF3*, *PTPRD*, *RAB31*, *RCBTB2*; Figure 3E), indicating coordinated changes in striatal gene networks implicated in neuronal stress associated with opioid addiction. Additional links included modules for NFATC2 and RXRG (FDR < 0.042; Figure 3E). Both nuclear factor of activated T-cell C2 (NFATC2) and the retinoid X receptor gamma (RXRG) of the RXR family of receptors link changes in neuronal activity associated with addictive drugs to processes involved in neuroprotection, neurodegeneration,^84–87^ and reward-related behaviors^51, 88–91^.

### Enrichment of DNA damage markers in specific neuronal subtypes in OUD

Several pathways and key transcription factor modules were related to processes of DNA damage and repair processes in OUD. Based on our results and other findings^9^, we investigated further the relationship between DNA damage^13^ and OUD in specific cell types (Figure 4A). In striatal neurons, we observed a significant enrichment of DNA damage markers in OUD subjects (p=0.046; linear regression t=2.18; Figure 4B). More specifically, a significant increase in DNA damage markers were found in interneurons of OUD subjects (p=0.035, linear regression t=2.14; Figure 4C, S6; Table S10). We further resolve augmented markers of DNA damage across specific striatal interneuron subtypes with the exception of *PTHLH*^+^ interneurons in OUD subjects (Figure S7, p < 0.025, linear regression). Our findings suggest striatal neurons may incur DNA damage in response to opioids and other factors, such as stress, neuroinflammation, and hypoxia (Figure 3), associated with opioid addiction. Interestingly, hypoxia pathways were significantly downregulated in OUD (Figure 3D; Table S9), including decreased expression of genes that are activated in response to hypoxic events (*e.g., NMRK1, BHLHE40;* Figure 3A)^92–94^, or overexpression of hypoxia-inducible transcription factor, HIF1A. Therefore, while respiratory depression is a hallmark of opioid overdose, downregulation of hypoxia-responsive genes points toward possible compensation in neurons secondary to periodic hypoxia and altered redox states, consistent with rodent models of substance use^95^.

**Figure 4.**
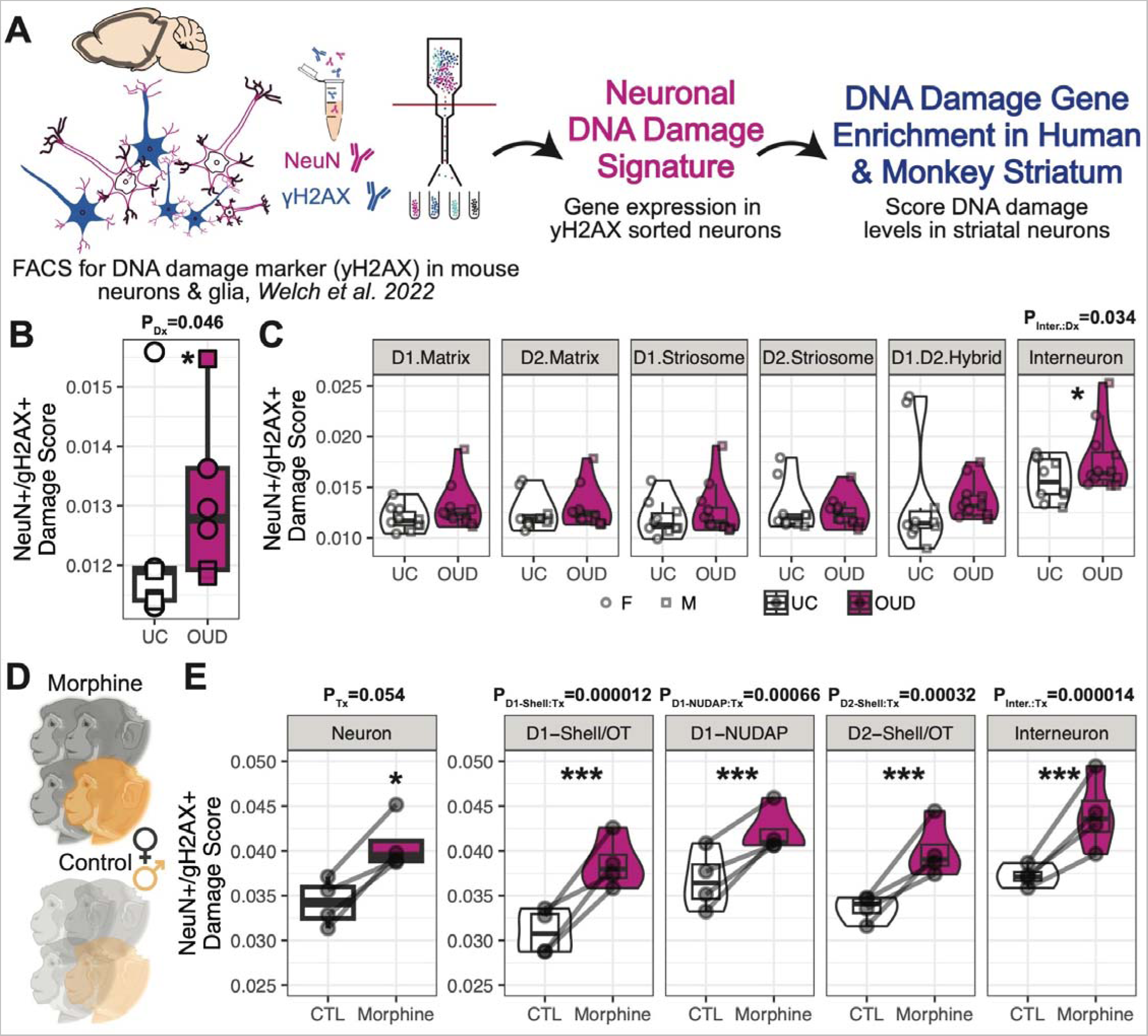
Elevated markers of DNA damage in striatal neurons associated with OUD and chronic morphine. (A) Schematic of the application of DNA damage gene signatures from a mouse model of Alzheimer’s Disease^13^ to score DNA damage in human striatal neurons. (B) Boxplot of subject-level pseudobulk average DNA damage scores across striatal neurons between unaffected and OUD subjects (P_Dx_ = 0.046, linear regression). (C) Violin-boxplot of cell type-level pseudobulk average DNA damage scores between unaffected and OUD subjects. The significant effect of diagnosis P-values from linear regressions are reported above each plot. (D) Schematic of chronic morphine exposure or unexposed rhesus macaques (N=4 subjects per treatment). (E) Boxplots of subject level pseudobulk average DNA damage scores across striatal neurons or neuronal subtypes between control subjects or those exposed to chronic morphine. The significant effect of chronic morphine treatment P-values from linear regressions are reported above each plot. (* P < 0.05; ** P < 0.01; *** P < 0.001).

Subjects with OUD were selected based on time since initial diagnosis of at least four years and other key factors, capturing the consequences of long-term opioid use on the human brain. OUD subjects also died of accidental opioid overdose, which leads to the possibility of introducing the impact of acute opioids and other substances on the brain. To further investigate the consequences of long-term opioid use on DNA damage-related processes in the brain, we assessed DNA damage marker enrichment in striatal neurons of non-human primates using snRNAseq (Figure 4D). Male and female rhesus macaques were administered morphine for ∼6 months, twice daily, then brain tissue was collected and striatum was rapidly dissected and stored (Figure 4D, S8A-B). Rhesus macaques were matched on age, sex, and body weight between morphine and control groups (Figure S8A). Cell types were clustered across rhesus macaque subjects, which were consistent with our previously described cell populations in non-human primate striatum (Figure S8C)^37^. Consistent with increased DNA damage in striatum of OUD subjects, rhesus macaques displayed significant enrichment of DNA damage markers in neurons in response to chronic opioid administration (P = 0.05, linear regression; Figure 3E). Significant enrichment was also identified across neuronal subtypes of opioid administered rhesus macaques (P < 0.00066, linear regression; Figure 3E), despite limitations (*i.e.,* species differences of striatal regions and species-specific opioid-induced gene expression changes) (Figure S8, S9)^96^. Thus, overall, elevated markers of DNA damage were evident in neurons of postmortem brains from subjects with OUD and non-human primates chronically administered opioids.

### Enriched pathways among glial cells support molecular signatures related to neuroinflammation and synaptic signaling in OUD

Compared to neurons, we identified more than twice as many DEGs in glial cells between unaffected and OUD subjects (Figure 2B, 4A). The number of DEGs were independent of cell type proportions and sequencing depth for these cell types, as microglia and endothelial cells had the most DEGs among glial cells, and expression patterns were largely distinct between glial cell types (Figure S6). An exception was the robust enrichment of genes involved in interferon response across multiple glial cell types, including astrocytes and oligodendrocytes (Figure 5C). Interferon response genes were significantly upregulated in all glial cell types in OUD subjects (FDR < 0.047, Table S11). Upregulated hub genes within the interferon response pathway included: *CMP2*, *HLA-F*, *IFI44*, *PPM1B*, and *RSAD2* (Figure 5A). Furthermore, we observed an upregulation of the NFKB1 transcription factor gene regulatory module in astrocytes (Figure 5B,E) and the upregulation of the STAT3 module in oligodendrocyte precursor cells (OPC,FDR< 0.036; Figure 5B). Several of the predicted NFKB1 gene targets were also differentially expressed including the inhibitor of this pathway, *NFKBIA* (log2FC = 1.86, FDR = 0.039; Figure 5E). Other glial cell types showed non-significant activation of this module, with the weakest being microglia. In response to cell stress, microglia release pro-inflammatory cytokines that may trigger an interferon response in other cell types, potentially responsible for the more broad patterns of interferon activation we observe across multiple cell types in human striatum associated with OUD^13, 97^.

**Figure 5.**
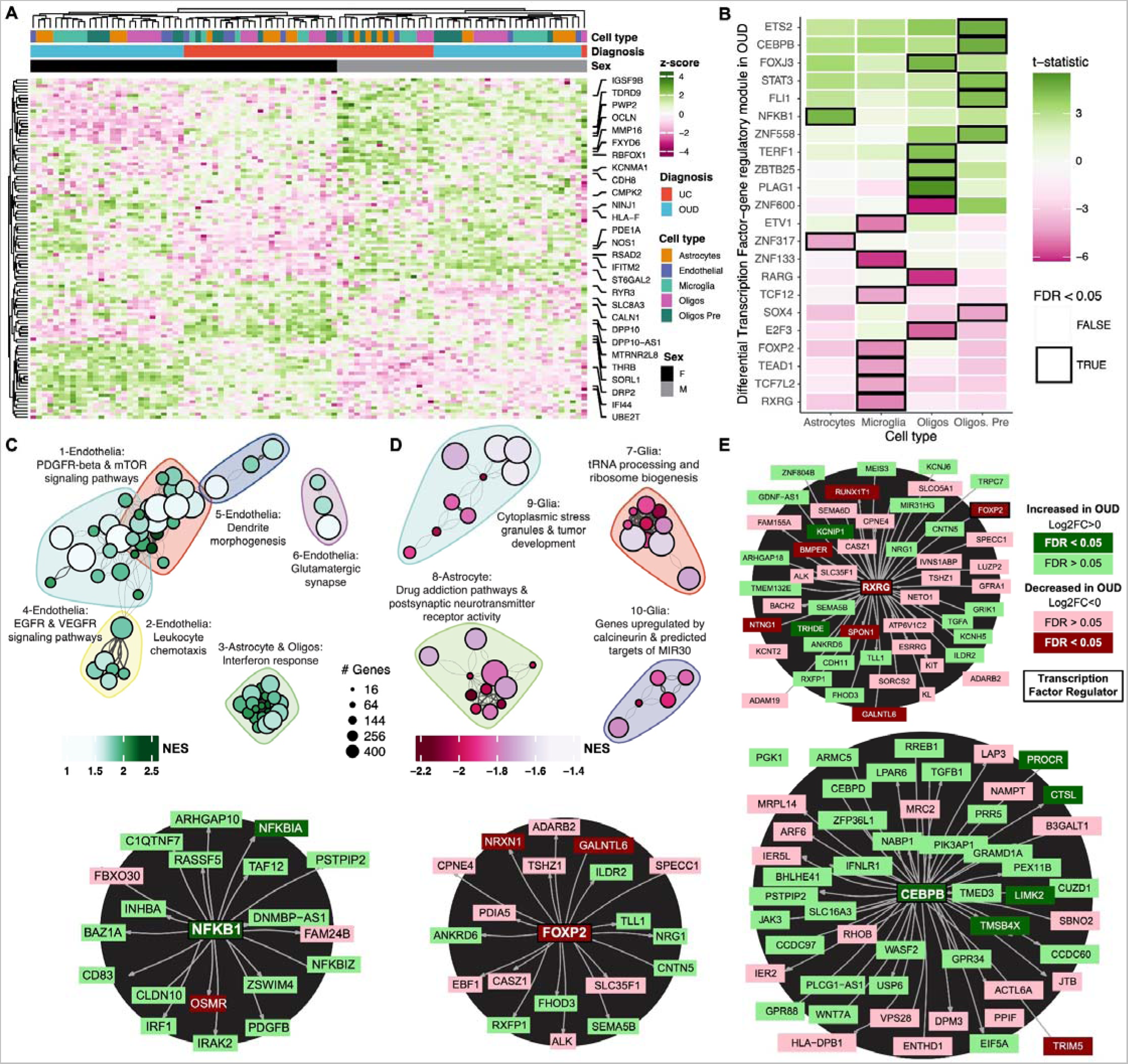
Transcriptional alterations in specific glial cell types in human striatum associated with OUD. (A) Expression heatmap of z-normalized pseudobulk gene expression, where the rows are DEGs and the columns are the biological replicates labeled by cell type, diagnosis, and sex of each subject. Genes that are represented by the enriched pathways in (C-D) are labeled along the right margins. (B) A heatmap plotting the t-statistic of differentially activated transcription factor gene regulatory networks across glial subtypes in OUD. (C-D) Network plot of significant clusters and enriched pathways, where each point is a significantly enriched pathway in a glial cell type and lines represent the proportion of shared genes between two pathways. Each point is colored by the normalized enrichment score (NES), indicating whether a pathway is enriched in up- or down-regulated genes. The color outlines represent the unique clusters of interconnected pathways by shared genes and the nearby text labels the unique cluster number and briefly summarizes which cell types and pathways are represented. (E) Cluster maps of select transcription factor gene regulatory networks. Each transcription factor or gene is colored to denote the direction and significance of DE in glial subtypes in OUD.

Altered neuroinflammatory signaling in microglia has previously been reported in postmortem brains from subjects with OUD^5^. Several DEGs in microglia lend further support roles of inflammation in synaptic plasticity in OUD^98^. For example, the PAX-FOXO1, CDH1, and neuron projection signaling pathways were enriched among microglia DEGs (FDR < 0.049). PAX-FOXO1 signaling regulates cellular senescence in the brain and is involved in the pathogenesis of several neurodegenerative disorders^15, 99^, while CDH1 signaling is involved in glial cell migration and axonal projections^100^. In the neuronal projection pathway, *ADGRB3* was among the top downregulated genes in microglia of OUD subjects (FDR = 3.5e-7). *ADGRB3* (known as BAI3, brain-specific angiogenesis inhibitor 3) encodes for a protein that has high-affinity for complement, C1q, an innate immune component released by neurons to eliminate damaged or inactive synapses^101–103^. Downregulation of *ADGRB3* in microglia may lead to aberrant synaptic formation.

In support of synaptic changes associated with opioid addiction, several genes involved in glutamatergic and GABAergic synaptic functions were altered in subjects with OUD (astrocytes: metabotropic glutamate receptor 5, *GRM5*, log2FC = -3.13, FDR = 1.06 E -14 and glutamate ionotropic receptor AMPA type subunit 1, *GRIA1*, log2FC = -1.98, FDR = 2.8 E -8; oligodendrocytes: GABA type A receptor subunit beta1, *GABRB1*, log2FC = -0.717, FDR = 0.023; microglia: GABA type A receptor subunit gamma 2, *GABRG2*, log2FC = -1.45, FDR = 0.038). Indeed, we also identified several pathways related to various synaptic (*e.g.,* glutamatergic synapse) and immune functions (*e.g.,* (TNF-alpha signaling) in gene co-expression network modules unique to MSN subpopulations and microglia (Figure S10-12; Table S18, S19). Together, our cell type-specific findings suggest an interplay between microglial-dependent signaling and synaptic plasticity in OUD, consistent with our previous bulk transcriptomics findings in other striatal subregions from subjects with OUD^5^. With single nuclei resolution, we report that the elevation of neuroinflammatory pathways related to microglial activation in OUD is likely due to transcriptional alterations within microglia, rather than pronounced changes in the number of microglia within striatum in subjects with OUD (FDR = 0.48, linear mixed effect regression for differential abundance; Figure 1F; Table S2).

Endothelial cells displayed the second most DEGs in OUD. Upregulated pathways in endothelial cells included several growth factors, VEGFR, EGFR, and PDGFR^104–106^ (Figure 5C). Several of these growth factors are linked to nociception and opioid tolerance^105, 107^. Other pathways included leukocyte chemotaxis and dendrite morphogenesis. Among the densely connected endothelial pathways, the top significantly upregulated hub gene was the brain-specific angiogenesis inhibitor 1-associated protein 2, *BAIAP2* (log2FC = 2.31, FDR = 0.0017), and, interestingly, the Alzheimer’s related microtubule-associated protein tau, *MAPT* (log2FC = 1.58, FDR = 0.036). The upregulation of *MAPT* may contribute to neurovascular brain insults in OUD^108, 109^.

The identification of differentially expressed transcription factor gene regulatory modules across glial cell types highlights several key factors in the regulation of immune response related to OUD. In oligodendrocyte precursor cells, the CEBPB module was upregulated in OUD (FDR = 0.031; Figure 5B,E), with prior studies showing diverse functions of CEBPB in proinflammatory states^110^ and the direct regulation of *APOE* in Alzheimer’s disease-related pathologies^111, 112^ . Notably, *APOE* is concordantly upregulated in oligodendrocyte precursor cells (log2FC = 2.56, FDR = 5.9e-4), along with other predicted CEBPB targets (Figure 5B,E). The RXRG transcription factor-gene regulatory module, which was downregulated in neuronal subtypes, was also downregulated in microglia (Figure 5B,E, FDR = 0.027). An inferred target of RXRG is *FOXP2* (Figure 5E), which was recently associated with OUD and other substance use disorders at the genome-wide level^113^. FOXP2 expression in microglia may be unique to humans compared to other primates^114^. Thus, downregulation of the FOXP2 transcription factor gene regulatory module in human microglia may link the genetic risk of OUD to other traits, including risk-taking behaviors^115^, and neurodevelopmental processes^116^.

### Sex-specific transcriptional alterations across striatal cell types associated with OUD

Prevalence rates of OUD and responses to opioids are dependent on sex^117–120^. Between unaffected and OUD subjects, we found a significant effect of sex on DEGs in both neuronal and glial cell types. Overall, the impact of sex on DEGs in OUD was magnified in neurons (Figure 3A) relative to glia (Figure 5A). Therefore, we conducted complementary, secondary analyses to identify sex-specific transcriptional alterations in neurons and glia associated with OUD (Table S12). First, we identified DEGs within either females or males in OUD subjects (Figure 6A, FDR < 0.05), revealing more sex-specific DEGs in glial cells relative to neurons. Additionally, we found a higher number of DEGs in females than males, 360 ± 60 versus 212 ± 67, respectively, in glial cells, and similarly in neurons (females: 91 ± 9.1; and males: 66 ± 7.9. Second, we explored the interaction between sex and diagnosis across genes and cell types to identify gene alterations occurring in only males or females of OUD subjects. Similar to our analysis of the main effect of sex in OUD, we found more DEGs in glial cells (291 ± 55) than neurons (97 ± 5.6) that have sex-specific changes in OUD (FDR < 0.05, Table S12). Among glial cell types, more DEGs were identified in females with OUD relative to males (Figure S13). The complete set of gene alterations associated with OUD and sex are reported in Table S12. Lastly, we identified DEGs that were different between females and males within diagnosis. As expected, sex-specific DEGs in OUD subjects compared to unaffected subjects were largely different, suggesting gene alterations dependent on sex and diagnosis were independent of naturally occurring gene expression variations between sexes (Figure S14). Collectively, our findings indicated that many genes altered in OUD depend on sex across striatal cell types, especially glial cells.

**Figure 6.**
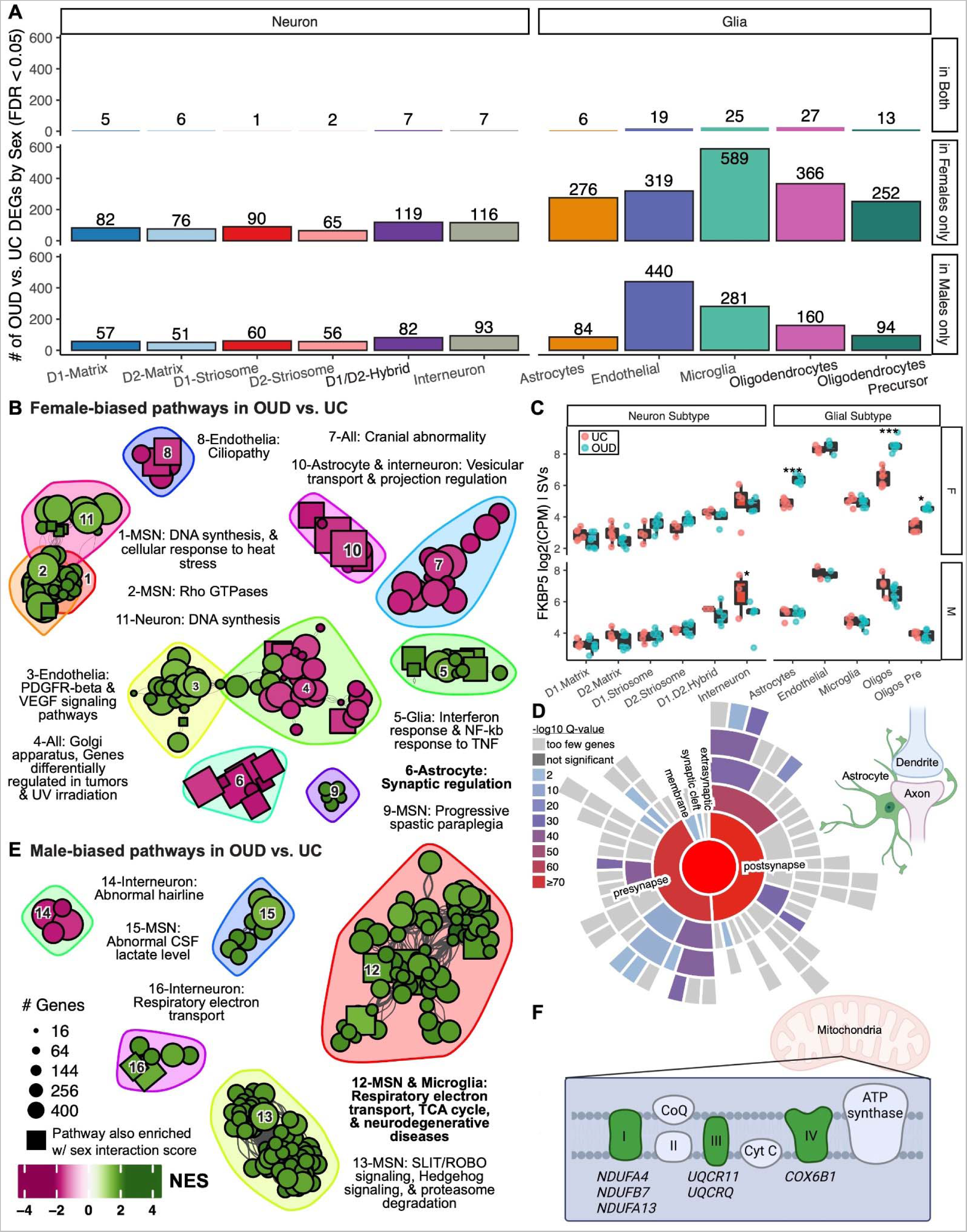
Sex-biased transcriptional alterations in striatal cell types associated with OUD. (A) Barplot showing the number of DEGs detected by cell type in sex-specific differential analyses comparing unaffected and OUD subjects. The bar plots show significant DEGs in only female subjects, in only male subjects, or in both groups (FDR < 0.05). (B) Network plot of significant clusters and enriched pathways as in Figure 2B-C and 4B-C. Here, enriched pathways include only DEGs calculated within female subjects if less significant than those calculated within male subjects. Pathways with a square point are also enriched using an alternate calculation of a sex-interaction score. (C) Boxplots showing sex- and cell type-specificity of differential expression of *FKBP5*. The boxplot labels the log2(counts per million) normalized gene expression values regressing out covariates and surrogate variables. Each point is a biological replicate colored by diagnosis (* P_Dx within Sex_ < 0.05; ** P_Dx within Sex_ < 0.01; *** P_Dx within Sex_ < 0.001). (D) A sunburst plot representing curated synaptic gene sets from SynGO from genes enriched in cluster 6, a female-biased set of pathways enriched in astrocytes. (E) A network plot as in (B); however, displaying the male-biased pathways. (F) Diagram showing the male-biased DEGs that are components of the electron transport gene.

To investigate which biological processes underlie sex-specific changes in OUD, we performed pathway analyses using female- and male-biased DEGs. In female OUD subjects, pathways were primarily associated with the upregulation of DNA repair processes, particularly in MSNs, accompanied by upregulation of interferon response signaling in glial cells, along with the upregulation of synaptic-related functions in astrocytes (Figure 6B; Table S13; Table S15). We then investigated significant key sex-specific factors in specific cell types of OUD subjects. We identified the FK506 binding protein 5, *FKBP5*, as a potential factor in the role of sex in opioid use. In the brain, FKBP5^121, 122^ is likely a key factor involved in stress and opioids in females^123^. *FKBP5* was significantly upregulated in astrocytes, oligodendrocytes, and oligodendrocyte precursor cells of female OUD subjects (Figure 6C; log2FC_F glia_>1.30, FDR_F glia_ < 0.0165, log2FC_M_= - 1.62, FDR_M interneurons_ < 0.040).

Preclinical rodent studies found marked sex differences of *Fkbp5* expression in dorsal striatum following opioid administration^124, 125^, and thus, investigated whether there were similarities in gene changes between rodents and humans in dorsal striatum. Indeed, gene sets from rodents exposed to opioids were significantly enriched in both glia and oligodendrocytes of females with OUD (FDR_F glia_ < 0.027; Table S16). Cross-species enriched DEGs were enriched for downregulation of synaptic regulation, particularly in astrocytes (Figure 6B; cluster 6), spanning both pre- and postsynaptic compartments (Figure 6D; Table S14; Table S15). In contrast, male-biased transcriptional alterations in OUD were mostly found in microglia, MSNs, and interneurons (Figure 6E; clusters 12, 15, 16), consisting of pathways related to mitochondrial functions (Figure 6F). Collectively, our evidence of sex differences in cell type-specific molecular signaling suggests an augmented, magnified glial cell response in females compared to males with OUD.

## Discussion

Using single nuclei transcriptomics technologies, we identified both canonical neuronal and glial cell types in human striatum (*e.g.,* D1 and D2 MSNs), along with several of the less abundant cell types, including subclasses of striatal interneurons and D1/D2-hybrid neurons. We also characterized the expression of specific opioid receptor and endogenous ligand subclasses across cell types, highlighting both expression patterns unique to human striatum. Our single nuclei findings provide further insights into the specific cell types that may be impacted in OUD, while highlighting several putative mechanisms in human striatal neuronal and glial subtypes. A common theme that emerged from our analyses was the involvement of processes related to neuroinflammation and cell stress in OUD, including a broad interferon response among glial cells and elevated DNA damage in neurons. DNA damage markers were identified in both the striatum of humans and monkeys, suggesting chronic opioid use associated with OUD leads to augmented DNA modifications (*e.g.,* double-stranded breaks, chromatin accessibility). Elevated markers for neuroinflammation, the involvement of microglia-dependent signaling in striatum, and alterations in synaptic signaling in OUD are consistent with previous bulk transcriptomics findings from postmortem brains of subjects with OUD^2, 34, 108, 126, 127^.

Many of the genes and pathways enriched in specific striatal cell types of OUD subjects have been implicated in various processes related to cell stress and senescence, DNA damage, and inflammation. Opioids lead to persistent changes in neuronal activity and synaptic plasticity within striatum. Prolonged or repeated alterations in neuronal activity combined with dysfunction within DNA repair pathways hampered by persistent cell stress (*e.g.,* oxidative stress) may lead to the accumulation of DNA damage. Neurons may be particularly vulnerable to DNA damage^128^. Integrity of DNA is critical for neural cells to prevent insertions, deletions, or mutations that can lead to aberrant synaptic plasticity and/or neurodegeneration. Double-stranded breaks and subsequent DNA repair may allow neurons to rapidly respond at the transcriptional level to changes in neuronal activity^129–132^. Our results suggest elevated DNA damage in neurons of OUD subjects, particularly within striatal interneurons.

Several subclasses of striatal interneurons are found in striatum^133^, possessing an array of transcriptional and physiological properties. We separately clustered TH+, CCK+, PTHLH+, and SST+ interneurons in human striatum. The functionality of interneurons in the striatum is diverse, with several subclasses modulating the excitability and inhibition of MSNs in response to various substances^134^, including opioids^135^. Although we were unable to reference the overall elevation of DNA damage signature to a specific subset of interneurons, our findings suggest striatal interneurons may be particularly vulnerable to a loss of DNA integrity and repair mechanisms in response to changes in neuronal activity by opioids and related perturbations.

The DNA damage response in neurons can lead to a proinflammatory response in microglia and other glial cells^13, 97^. In OUD, there were a relatively large number of genes that were upregulated specifically in microglia, enriched for genes involved in neuroinflammation. Microglial activation may propagate to other neural cell types, evident by an increase in interferon responses^13, 97^. Indeed, we found significantly upregulated interferon response in both neuronal and glial cells of OUD subjects. Interferon responses may modulate opioid actions in the brain and could be involved in the emergence of symptoms of opioid withdrawal^136^.

Other relevant pathways to OUD included oxidative stress, mitochondrial respiration, and neuroprotection. The coordinated alterations in the expression of genes involved in these pathways may reflect compensation to changes in neural activity impacted by opioid use. For example, glutamatergic hyperactivity at striatal MSNs^137^ may cascade towards excitotoxicity and elevated DNA damage^14, 23, 138^, both of which have been recently associated with OUD^139, 140^. GRM5 was specifically downregulated in astrocytes in OUD. GRM5 is transiently expressed in astrocytes to detect and respond to extracellular glutamate, suggesting downregulation in OUD may be due to changes in glutamatergic activity in striatal neurons^141^. Alterations in other glutamatergic and GABAergic transcripts were also found in specific cell types in OUD (*e.g.,* downregulation of *GRIA1* in astrocytes and upregulation of *GABRG2* in microglia). In parallel, enrichment of neurodegenerative-related and neuronal activity markers in certain striatal cells in OUD may be a result of high energetic demand on MSNs to maintain a hyperpolarized state, a possible mechanism of vulnerability of striatal MSNs proposed in Huntington’s disease^142^. We also found OUD-associated changes in neuronal energetics with the downregulation of nicotinamide riboside kinase 1, *NMRK1*, and the basic helix-loop-helix family member e40, *BHLHE40* (DEC1), both of which are involved in regulating cellular metabolism, oxidative stress, and inflammation^92–94^. Together, the broad set of changes in genes and pathways associated with OUD were associated with various processes involved in cell stress.

Sex differences in the vulnerability to substance use, development of substance use disorders, and treatment outcomes, along with the biological response to drugs are evident from both preclinical and clinical studies^143–145^. Our findings suggest sex-specific alterations in the transcriptional response to opioids in specific striatal cell types. Based on the transcriptional patterns, we found a more pronounced inflammatory response associated with striatal microglia in females compared to males with OUD, suggesting microglia in the human brain also exhibit sex-specific responses to stress and substances^145–147^. In addition, we identified the upregulation of *FKBP5* as a potential female-specific factor in several glial subtypes in OUD. *FKBP5* acts as a co-chaperone of the glucocorticoid receptor activated in response to stress, with major implications in the pathology of several psychiatric disorders and the impact of stress on substance withdrawal, craving, and relapse^148–150^. Further, the transcriptional alterations within microglia, oligodendrocytes, and astrocytes in female OUD subjects were significantly enriched for *FKBP5* target genes identified in rodents administered opioids. Genes related to synaptic functions were also enriched in astrocytes of females, while enrichment was mostly in MSNs and interneurons in male OUD subjects, suggesting sex-specific alterations of striatal neuronal and glial cell signaling in opioid addiction.

Several limitations of our resource stem from challenges of profiling single cell transcriptomes in postmortem human brains. While the nuclear transcriptome correlates on whole with the transcriptomes of other subcellular compartments, there are notable differences in mRNA trafficking within certain neural cell types and synapses. Furthermore, the nuclear transcriptome does not capture post-transcriptional regulation. Further, the sex-biased differential expression in OUD is likely influenced by cultural and behavioral factors that are confounded with biological sex. Lastly, while this resource will be useful to identify the consequences of OUD in the postmortem brain transcriptome, integration of our findings with other types of approaches from genetics to functional genomics in clinical cohorts to animal models of opioid addiction could contribute in a greater understanding of opioid addiction and aid in the realization of new treatment strategies.

## Supporting information

Supplemental Methods and Results

## References

1. National Institute on Drug Abuse. Drug Overdose Death Rates. National Institute on Drug Abuse https://nida.nih.gov/research-topics/trends-statistics/overdose-death-rates (2023).

2. Puig, S., et al. Uncovering circadian rhythm disruptions of synaptic proteome signaling in prefrontal cortex and nucleus accumbens associated with opioid use disorder. bioRxiv (2023) doi:10.1101/2023.04.07.536056.

3. Nagamatsu, S. T. et al. CpH methylome analysis in human cortical neurons identifies novel gene pathways and drug targets for opioid use disorder. Front. Psychiatry 13, 1078894 (2022).

4. Xue, X. et al. Molecular rhythm alterations in prefrontal cortex and nucleus accumbens associated with opioid use disorder. Transl. Psychiatry 12, 123 (2022).

5. Seney, M. L. et al. Transcriptional Alterations in Dorsolateral Prefrontal Cortex and Nucleus Accumbens Implicate Neuroinflammation and Synaptic Remodeling in Opioid Use Disorder. Biol. Psychiatry 90, (2021).

6. Mendez, E. F. et al. Angiogenic gene networks are dysregulated in opioid use disorder: evidence from multi-omics and imaging of postmortem human brain. Mol. Psychiatry 26, 7803–7812 (2021).

7. Egervari, G. et al. Chromatin accessibility mapping of the striatum identifies tyrosine kinase FYN as a therapeutic target for heroin use disorder. Nat. Commun. 11, 4634 (2020).

8. Kovacs, G. G. et al. Heroin abuse exaggerates age-related deposition of hyperphosphorylated tau and p62-positive inclusions. Neurobiol. Aging 36, 3100–3107 (2015).

9. Wang, Y. et al. Opioid induces increased DNA damage in prefrontal cortex and nucleus accumbens. Pharmacol. Biochem. Behav. 224, 173535 (2023).

10. Zahmatkesh, M., Kadkhodaee, M., Salarian, A., Seifi, B. & Adeli, S. Impact of opioids on oxidative status and related signaling pathways: An integrated view. J. Opioid Manag. 13, 241–251 (2017).

11. Calarco, C. A. et al. Mitochondria-Related Nuclear Gene Expression in the Nucleus Accumbens and Blood Mitochondrial Copy Number After Developmental Fentanyl Exposure in Adolescent Male and Female C57BL/6 Mice. Front. Psychiatry 12, 737389 (2021).

12. Song, X. et al. DNA Repair Inhibition Leads to Active Export of Repetitive Sequences to the Cytoplasm Triggering an Inflammatory Response. J. Neurosci. 41, 9286–9307 (2021).

13. Welch, G. M. et al. Neurons burdened by DNA double-strand breaks incite microglia activation through antiviral-like signaling in neurodegeneration. Sci Adv 8, eabo4662 (2022).

14. Martin, L. J. & Chang, Q. DNA Damage Response and Repair, DNA Methylation, and Cell Death in Human Neurons and Experimental Animal Neurons Are Different. J. Neuropathol. Exp. Neurol. 77, 636–655 (2018).

15. Liu, L. et al. Cross-Talking Pathways of Forkhead Box O1 (FOXO1) Are Involved in the Pathogenesis of Alzheimer’s Disease and Huntington’s Disease. Oxid. Med. Cell. Longev. 2022, (2022).

16. Lawler, A. J. et al. Cell Type-Specific Oxidative Stress Genomic Signatures in the Globus Pallidus of Dopamine-Depleted Mice. J. Neurosci. 40, 9772–9783 (2020).

17. Araújo, B. et al. Neuroinflammation and Parkinson’s disease-from neurodegeneration to therapeutic opportunities. Cells 11, 2908 (2022).

18. Becher, B., Spath, S. & Goverman, J. Cytokine networks in neuroinflammation. Nat. Rev. Immunol. 17, 49–59 (2017).

19. Avey, D. et al. Single-Cell RNA-Seq Uncovers a Robust Transcriptional Response to Morphine by Glia. Cell Rep. 24, 3619–3629.e4 (2018).

20. Madabhushi, R., Pan, L. & Tsai, L.-H. DNA damage and its links to neurodegeneration. Neuron 83, 266–282 (2014).

21. Shanbhag, N. M. et al. Early neuronal accumulation of DNA double strand breaks in Alzheimer’s disease. Acta Neuropathologica Communications vol. 7 Preprint at 10.1186/s40478-019-0723-5 (2019).

22. Wang, Z.-X., Li, Y.-L., Pu, J.-L. & Zhang, B.-R. DNA damage-mediated neurotoxicity in Parkinson’s disease. Int. J. Mol. Sci. 24, 6313 (2023).

23. Yang, J.-L., Sykora, P., Wilson, D. M., 3rd, Mattson, M. P. & Bohr, V. A. The excitatory neurotransmitter glutamate stimulates DNA repair to increase neuronal resiliency. Mech. Ageing Dev. 132, 405–411 (2011).

24. Mathys, H. et al. Single-cell transcriptomic analysis of Alzheimer’s disease. Nature 570, 332–337 (2019).

25. Geloso, M. C. et al. The Dual Role of Microglia in ALS: Mechanisms and Therapeutic Approaches. Front. Aging Neurosci. 9, 242 (2017).

26. Shokri-Kojori, E. et al. Brain opioid segments and striatal patterns of dopamine release induced by naloxone and morphine. Hum. Brain Mapp. 43, (2022).

27. Shokri-Kojori, E., Wang, G.-J. & Volkow, N. D. Naloxone precipitated withdrawal increases dopamine release in the dorsal striatum of opioid dependent men. Transl. Psychiatry 11, 445 (2021).

28. Koob, G. F. & Volkow, N. D. Neurobiology of addiction: a neurocircuitry analysis. The lancet. Psychiatry 3, 760 (2016).

29. Zilverstand, A., Huang, A. S., Alia-Klein, N. & Goldstein, R. Z. Neuroimaging Impaired Response Inhibition and Salience Attribution in Human Drug Addiction: A Systematic Review. Neuron 98, (2018).

30. Matsushima, A. et al. Transcriptional vulnerabilities of striatal neurons in human and rodent models of Huntington’s disease. Nat. Commun. 14, 282 (2023).

31. Reiss, D., Maduna, T., Maurin, H., Audouard, E. & Gaveriaux-Ruff, C. Mu opioid receptor in microglia contributes to morphine analgesic tolerance, hyperalgesia, and withdrawal in mice. J. Neurosci. Res. 100, 203–219 (2022).

32. Zhang, H., Largent-Milnes, T. M. & Vanderah, T. W. Glial neuroimmune signaling in opioid reward. Brain Res. Bull. 155, 102–111 (2020).

33. Bachtell, R. K., Jones, J. D., Heinzerling, K. G., Beardsley, P. M. & Comer, S. D. Glial and neuroinflammatory targets for treating substance use disorders. Drug Alcohol Depend. 180, 156–170 (2017).

34. Seney, M. L. et al. Transcriptional Alterations in Dorsolateral Prefrontal Cortex and Nucleus Accumbens Implicate Neuroinflammation and Synaptic Remodeling in Opioid Use Disorder. Biol. Psychiatry 90, (2021).

35. Coffey, K. R. et al. A cAMP-Related Gene Network in Microglia Is Inversely Regulated by Morphine Tolerance and Withdrawal. Biol Psychiatry Glob Open Sci 2, 180–189 (2022).

36. Mogali, S., Askalsky, P., Madera, G., Jones, J. D. & Comer, S. D. Minocycline attenuates oxycodone-induced positive subjective responses in non-dependent, recreational opioid users. Pharmacol. Biochem. Behav. 209, 173241 (2021).

37. He, J. et al. Transcriptional and anatomical diversity of medium spiny neurons in the primate striatum. Curr. Biol. 31, 5473–5486.e6 (2021).

38. Gokce, O. et al. Cellular Taxonomy of the Mouse Striatum as Revealed by Single-Cell RNA-Seq. Cell Rep. 16, 1126–1137 (2016).

39. Stanley, G., Gokce, O., Malenka, R. C., Südhof, T. C. & Quake, S. R. Continuous and discrete neuron types of the adult murine striatum. Neuron 105, 688–699.e8 (2020).

40. Saunders, A. et al. Molecular Diversity and Specializations among the Cells of the Adult Mouse Brain. Cell 174, (2018).

41. Krienen, F. M. et al. Innovations present in the primate interneuron repertoire. Nature 586, 262–269 (2020).

42. Tran, M. N. et al. Single-nucleus transcriptome analysis reveals cell-type-specific molecular signatures across reward circuitry in the human brain. Neuron 109, 3088–3103.e5 (2021).

43. 43. Gayden, J., et al. Three-dimensional characterization of medium spiny neuron heterogeneity in the adult mouse striatum. *bioRxiv* (2023) doi:10.1101/2023.05.04.539488.

44. Sharif, N. A. & Hughes, J. Discrete mapping of brain Mu and delta opioid receptors using selective peptides: quantitative autoradiography, species differences and comparison with kappa receptors. Peptides 10, 499–522 (1989).

45. Maduna, T. et al. Microglia Express Mu Opioid Receptor: Insights From Transcriptomics and Fluorescent Reporter Mice. Front. Psychiatry 9, 726 (2018).

46. Dai, K. Z. et al. Dopamine D2 receptors bidirectionally regulate striatal enkephalin expression: Implications for cocaine reward. Cell Rep. 40, 111440 (2022).

47. Crittenden, J. R. et al. Striosome-dendron bouquets highlight a unique striatonigral circuit targeting dopamine-containing neurons. Proc. Natl. Acad. Sci. U. S. A. 113, 11318–11323 (2016).

48. Hong, S. et al. Predominant Striatal Input to the Lateral Habenula in Macaques Comes from Striosomes. Curr. Biol. 29, (2019).

49. Aibar, S. et al. SCENIC: single-cell regulatory network inference and clustering. Nat. Methods 14, 1083–1086 (2017).

50. Samad, T. A., Krezel, W., Chambon, P. & Borrelli, E. Regulation of dopaminergic pathways by retinoids: activation of the D2 receptor promoter by members of the retinoic acid receptor-retinoid X receptor family. Proc. Natl. Acad. Sci. U. S. A. 94, 14349–14354 (1997).

51. Godino, A. et al. Transcriptional control of nucleus accumbens neuronal excitability by retinoid X receptor alpha tunes sensitivity to drug rewards. Neuron 111, 1453–1467.e7 (2023).

52. Mozzi, A. et al. A common genetic variant in FOXP2 is associated with language-based learning (dis)abilities: Evidence from two Italian independent samples. Am. J. Med. Genet. B Neuropsychiatr. Genet. 174, 578–586 (2017).

53. Deak, J. D. & Johnson, E. C. Genetics of substance use disorders: a review. Psychol. Med. 51, 2189–2200 (2021).

54. Sun, W. et al. SOX9 Is an Astrocyte-Specific Nuclear Marker in the Adult Brain Outside the Neurogenic Regions. J. Neurosci. 37, 4493–4507 (2017).

55. Mateusz, S. L. et al. β-catenin signaling via astrocyte-encoded TCF7L2 regulates neuronal excitability and social behavior. bioRxiv 2020.11.28.402099 (2020) doi:10.1101/2020.11.28.402099.

56. Matuzelski, E. et al. Transcriptional regulation of Nfix by NFIB drives astrocytic maturation within the developing spinal cord. Dev. Biol. 432, 286–297 (2017).

57. Kim, D. & Tsai, L.-H. Linking cell cycle reentry and DNA damage in neurodegeneration. Ann. N. Y. Acad. Sci. 1170, 674–679 (2009).

58. Nishimura, K. et al. Mcm8 and Mcm9 form a complex that functions in homologous recombination repair induced by DNA interstrand crosslinks. Mol. Cell 47, (2012).

59. Lee, K. Y. et al. MCM8-9 complex promotes resection of double-strand break ends by MRE11-RAD50-NBS1 complex. Nat. Commun. 6, 7744 (2015).

60. Osterweil, E., Wells, D. G. & Mooseker, M. S. A role for myosin VI in postsynaptic structure and glutamate receptor endocytosis. J. Cell Biol. 168, 329–338 (2005).

61. Auer, M., Hausott, B. & Klimaschewski, L. Rho GTPases as regulators of morphological neuroplasticity. Ann. Anat. 193, 259–266 (2011).

62. Tolias, K. F., Duman, J. G. & Um, K. Control of synapse development and plasticity by Rho GTPase regulatory proteins. Prog. Neurobiol. 94, 133–148 (2011).

63. Fritz, G. & Henninger, C. Rho GTPases: Novel Players in the Regulation of the DNA Damage Response? Biomolecules 5, 2417 (2015).

64. Ayton, S., Faux, N. G. & Bush, A. I. Ferritin levels in the cerebrospinal fluid predict Alzheimer’s disease outcomes and are regulated by APOE. Nat. Commun. 6, 1–9 (2015).

65. Reichert, C. O. et al. Ferroptosis Mechanisms Involved in Neurodegenerative Diseases. Int. J. Mol. Sci. 21, (2020).

66. Belaidi, A. A. et al. Apolipoprotein E potently inhibits ferroptosis by blocking ferritinophagy. Mol. Psychiatry 1–10 (2022).

67. Murphy, M. P. How mitochondria produce reactive oxygen species. Biochem. J 417, 1–13 (2009).

68. Shao, A.-W. et al. Bclaf1 is an important NF-κB signaling transducer and C/EBPβ regulator in DNA damage-induced senescence. Cell Death Differ. 23, 865–875 (2016).

69. Yu, Z., Zhu, J., Wang, H., Li, H. & Jin, X. Function of BCLAF1 in human disease. Oncol. Lett. 23, (2022).

70. McGowan, P. O. et al. Epigenetic regulation of the glucocorticoid receptor in human brain associates with childhood abuse. Nat. Neurosci. 12, 342–348 (2009).

71. Holmes, L., Jr et al. Aberrant Epigenomic Modulation of Glucocorticoid Receptor Gene (NR3C1) in Early Life Stress and Major Depressive Disorder Correlation: Systematic Review and Quantitative Evidence Synthesis. Int. J. Environ. Res. Public Health 16, (2019).

72. 72. Glucocorticoid receptor gene (NR3C1) methylation processes as mediators of early adversity in stress-related disorders causality: A critical review. Neurosci. Biobehav. Rev. 55, 520–535 (2015).

73. Green, T. A. et al. Induction of Activating Transcription Factors (ATFs) ATF2, ATF3, and ATF4 in the Nucleus Accumbens and Their Regulation of Emotional Behavior. J. Neurosci. 28, 2025–2032 (2008).

74. Pattinson, C. L. et al. Excessive daytime sleepiness is associated with altered gene expression in military personnel and veterans with posttraumatic stress disorder: an RNA sequencing study. Sleep 43, (2020).

75. Shahabi, N. A., McAllen, K. & Sharp, B. M. Phosphorylation of activating transcription factor in murine splenocytes through delta opioid receptors. Cell. Immunol. 221, (2003).

76. Banghart, M. R., Neufeld, S. Q., Wong, N. C. & Sabatini, B. L. Enkephalin Disinhibits Mu Opioid Receptor-Rich Striatal Patches via Delta Opioid Receptors. Neuron 88, 1227–1239 (2015).

77. Jiang, Z. G. & North, R. A. Pre- and postsynaptic inhibition by opioids in rat striatum. J. Neurosci. 12, 356–361 (1992).

78. Tajima, H. et al. Evidence for in vivo production of Humanin peptide, a neuroprotective factor against Alzheimer’s disease-related insults. Neurosci. Lett. 324, (2002).

79. Gurunathan, S., Jeyaraj, M., Kang, M.-H. & Kim, J.-H. Mitochondrial Peptide Humanin Protects Silver Nanoparticles-Induced Neurotoxicity in Human Neuroblastoma Cancer Cells (SH-SY5Y). Int. J. Mol. Sci. 20, (2019).

80. Ahuja, M. et al. Bach1 derepression is neuroprotective in a mouse model of Parkinson’s disease. Proc. Natl. Acad. Sci. U. S. A. 118, (2021).

81. Tolve, M. et al. The transcription factor BCL11A defines distinct subsets of midbrain dopaminergic neurons. Cell Rep. 36, (2021).

82. Simon, R., Wiegreffe, C. & Britsch, S. Bcl11 Transcription Factors Regulate Cortical Development and Function. Front. Mol. Neurosci. 13, (2020).

83. Cirnaru, M.-D. et al. Unbiased identification of novel transcription factors in striatal compartmentation and striosome maturation. Elife 10, (2021).

84. Benedito, A. B. et al. The transcription factor NFAT3 mediates neuronal survival. J. Biol. Chem. 280, 2818–2825 (2005).

85. Hudry, E. et al. Inhibition of the NFAT pathway alleviates amyloid β neurotoxicity in a mouse model of Alzheimer’s disease. J. Neurosci. 32, (2012).

86. Manocha, G. D. et al. NFATc2 Modulates Microglial Activation in the AβPP/PS1 Mouse Model of Alzheimer’s Disease. J. Alzheimers. Dis. 58, (2017).

87. Vihma, H., Luhakooder, M., Pruunsild, P. & Timmusk, T. Regulation of different human NFAT isoforms by neuronal activity. J. Neurochem. 137, (2016).

88. Krezel, W. et al. Impaired locomotion and dopamine signaling in retinoid receptor mutant mice. Science 279, 863– 867 (1998).

89. Krzyzosiak, A. et al. Retinoid x receptor gamma control of affective behaviors involves dopaminergic signaling in mice. Neuron 66, 908–920 (2010).

90. Wietrzych-Schindler, M. et al. Retinoid x receptor gamma is implicated in docosahexaenoic acid modulation of despair behaviors and working memory in mice. Biol. Psychiatry 69, 788–794 (2011).

91. Krzyżosiak, A. et al. Vitamin A5/X controls stress-adaptation and prevents depressive-like behaviors in a mouse model of chronic stress. Neurobiology of stress 15, (2021).

92. Cook, M. E., Jarjour, N. N., Lin, C.-C. & Edelson, B. T. Transcription Factor Bhlhe40 in Immunity and Autoimmunity. Trends Immunol. 41, 1023–1036 (2020).

93. Lin, C.-C. et al. Bhlhe40 controls cytokine production by T cells and is essential for pathogenicity in autoimmune neuroinflammation. Nat. Commun. 5, 1–13 (2014).

94. Zafar, A., Ng, H. P., Kim, G.-D., Chan, E. R. & Mahabeleshwar, G. H. BHLHE40 promotes macrophage pro-inflammatory gene expression and functions. FASEB J. 35, e21940 (2021).

95. Logan, R. W. et al. NAD+ cellular redox and SIRT1 regulate the diurnal rhythms of tyrosine hydroxylase and conditioned cocaine reward. Mol. Psychiatry 24, 1668–1684 (2019).

96. Xia, X. et al. Interspecies Differences in the Connectivity of Ventral Striatal Components Between Humans and Macaques. Front. Neurosci. 13, 623 (2019).

97. Roy, E. R. et al. Type I interferon response drives neuroinflammation and synapse loss in Alzheimer disease. J. Clin. Invest. 130, 1912–1930 (2020).

98. Butelman, E. R. et al. Neuroimmune mechanisms of opioid use disorder and recovery: Translatability to human studies, and future research directions. Neuroscience (2023) doi:10.1016/j.neuroscience.2023.07.031.

99. Thompson, J. A. & Ziman, M. Pax genes during neural development and their potential role in neuroregeneration. Prog. Neurobiol. 95, (2011).

100. Silies, M. & Klämbt, C. APC/CFzr/Cdh1-dependent regulation of cell adhesion controls glial migration in the Drosophila PNS. Nat. Neurosci. 13, 1357–1364 (2010).

101. Bolliger, M. F., Martinelli, D. C. & Südhof, T. C. The cell-adhesion G protein-coupled receptor BAI3 is a high-affinity receptor for C1q-like proteins. Proc. Natl. Acad. Sci. U. S. A. 108, 2534–2539 (2011).

102. Kakegawa, W. et al. Anterograde C1ql1 signaling is required in order to determine and maintain a single-winner climbing fiber in the mouse cerebellum. Neuron 85, 316–329 (2015).

103. Neniskyte, U. & Gross, C. T. Errant gardeners: glial-cell-dependent synaptic pruning and neurodevelopmental disorders. Nat. Rev. Neurosci. 18, 658–670 (2017).

104. Gamble, M. C. et al. Mu-opioid receptor and receptor tyrosine kinase crosstalk: Implications in mechanisms of opioid tolerance, reduced analgesia to neuropathic pain, dependence, and reward. Front. Syst. Neurosci. 16, (2022).

105. Puig, S., Donica, C. L. & Gutstein, H. B. EGFR Signaling Causes Morphine Tolerance and Mechanical Sensitization in Rats. eNeuro 7, (2020).

106. Barkai, O. et al. Platelet-derived growth factor activates nociceptive neurons by inhibiting M-current and contributes to inflammatory pain. Pain 160, (2019).

107. Lopez-Bellido, R. et al. Growth Factor Signaling Regulates Mechanical Nociception in Flies and Vertebrates. J. Neurosci. 39, 6012–6030 (2019).

108. Mendez, E. F. et al. Angiogenic gene networks are dysregulated in opioid use disorder: evidence from multi-omics and imaging of postmortem human brain. Mol. Psychiatry 26, 7803–7812 (2021).

109. Canepa, E. & Fossati, S. Impact of Tau on Neurovascular Pathology in Alzheimer’s Disease. Front. Neurol. 11, 573324 (2020).

110. Ko, C.-Y., Chang, W.-C. & Wang, J.-M. Biological roles of CCAAT/Enhancer-binding protein delta during inflammation. J. Biomed. Sci. 22, 6 (2015).

111. Xia, Y. et al. C/EBPβ is a key transcription factor for APOE and preferentially mediates ApoE4 expression in Alzheimer’s disease. Mol. Psychiatry 26, 6002–6022 (2020).

112. Yao, Y. et al. A delta-secretase-truncated APP fragment activates CEBPB, mediating Alzheimer’s disease pathologies. Brain 144, (2021).

113. Deak, J. et al. Genome-wide association study in individuals of European and African ancestry and multi-trait analysis of opioid use disorder identifies 19 independent genome-wide significant risk loci. Mol. Psychiatry 27, 3970–3979 (2022).

114. Ma, S. et al. Molecular and cellular evolution of the primate dorsolateral prefrontal cortex. Science 377, eabo7257 (2022).

115. Clifton, E. A. D. et al. Genome-wide association study for risk taking propensity indicates shared pathways with body mass index. Commun Biol 1, 36 (2018).

116. den Hoed, J., Devaraju, K. & Fisher, S. E. Molecular networks of the FOXP2 transcription factor in the brain. EMBO Rep. 22, e52803 (2021).

117. Gerdle, B. et al. Prevalence of widespread pain and associations with work status: a population study. BMC Musculoskelet. Disord. 9, (2008).

118. Ailes, E. C. et al. Opioid prescription claims among women of reproductive age--United States, 2008-2012. MMWR Morb. Mortal. Wkly. Rep. 64, (2015).

119. McHugh, R. K. et al. Gender differences in a clinical trial for prescription opioid dependence. J. Subst. Abuse Treat. 45, (2013).

120. Gjersing, L. & Bretteville-Jensen, A. L. Gender differences in mortality and risk factors in a 13-year cohort study of street-recruited injecting drug users. BMC Public Health 14, (2014).

121. Lee, R. S. et al. DNA methylation and sex-specific expression of FKBP5 as correlates of one-month bedtime cortisol levels in healthy individuals. Psychoneuroendocrinology 97, (2018).

122. Nold, V. et al. Impact of Fkbp5 × early life adversity × sex in humanised mice on multidimensional stress responses and circadian rhythmicity. Mol. Psychiatry 27, 3544–3555 (2022).

123. Serdarevic, M., Striley, C. W. & Cottler, L. B. Gender differences in prescription opioid use. Curr. Opin. Psychiatry 30, 238 (2017).

124. Piechota, M. et al. The dissection of transcriptional modules regulated by various drugs of abuse in the mouse striatum. Genome Biol. 11, R48 (2010).

125. McClung, C. A., Nestler, E. J. & Zachariou, V. Regulation of Gene Expression by Chronic Morphine and Morphine Withdrawal in the Locus Ceruleus and Ventral Tegmental Area. J. Neurosci. 25, 6005 (2005).

126. Xue, X. et al. Molecular rhythm alterations in prefrontal cortex and nucleus accumbens associated with opioid use disorder. Transl. Psychiatry 12, 123 (2022).

127. Mendez, E. F. et al. A human stem cell-derived neuronal model of morphine exposure reflects brain dysregulation in opioid use disorder: Transcriptomic and epigenetic characterization of postmortem-derived iPSC neurons. Front. Psychiatry 14, 1070556 (2023).

128. Wu, W. et al. Neuronal enhancers are hotspots for DNA single-strand break repair. Nature 593, 440–444 (2021).

129. Caldecott, K. W., Ward, M. E. & Nussenzweig, A. The threat of programmed DNA damage to neuronal genome integrity and plasticity. Nat. Genet. 54, 115–120 (2022).

130. Madabhushi, R. et al. Activity-Induced DNA Breaks Govern the Expression of Neuronal Early-Response Genes. Cell 161, 1592–1605 (2015).

131. Stott, R. T., Kritsky, O. & Tsai, L.-H. Profiling DNA break sites and transcriptional changes in response to contextual fear learning. PLoS One 16, e0249691 (2021).

132. Pollina, E. A. et al. A NPAS4-NuA4 complex couples synaptic activity to DNA repair. Nature 614, 732–741 (2023).

133. Muñoz-Manchado, A. B. et al. Diversity of Interneurons in the Dorsal Striatum Revealed by Single-Cell RNA Sequencing and PatchSeq. Cell Rep. 24, 2179–2190.e7 (2018).

134. Tepper, J. M. et al. Heterogeneity and Diversity of Striatal GABAergic Interneurons: Update 2018. Front. Neuroanat. 12, 91 (2018).

135. Arttamangkul, S., Platt, E. J., Carroll, J. & Farrens, D. Functional independence of endogenous μ- and δ-opioid receptors co-expressed in cholinergic interneurons. Elife 10, (2021).

136. Zhu, Y. et al. Opioid-induced fragile-like regulatory T cells contribute to withdrawal. Cell 186, 591–606.e23 (2023).

137. Koob, G. F. Drug Addiction: Hyperkatifeia/Negative Reinforcement as a Framework for Medications Development. Pharmacol. Rev. 73, 163–201 (2021).

138. Didier, M. et al. DNA strand breaks induced by sustained glutamate excitotoxicity in primary neuronal cultures. J. Neurosci. 16, 2238–2250 (1996).

139. Hearing, M. C. et al. Reversal of morphine-induced cell-type-specific synaptic plasticity in the nucleus accumbens shell blocks reinstatement. Proc. Natl. Acad. Sci. U. S. A. 113, 757–762 (2016).

140. Hearing, M., Graziane, N., Dong, Y. & Thomas, M. J. Opioid and Psychostimulant Plasticity: Targeting Overlap in Nucleus Accumbens Glutamate Signaling. Trends Pharmacol. Sci. 39, 276–294 (2018).

141. Panatier, A. & Robitaille, R. Astrocytic mGluR5 and the tripartite synapse. Neuroscience 323, 29–34 (2016).

142. Morigaki, R. & Goto, S. Striatal Vulnerability in Huntington’s Disease: Neuroprotection Versus Neurotoxicity. Brain Sci 7, (2017).

143. Lynch, W. J., Roth, M. E. & Carroll, M. E. Biological basis of sex differences in drug abuse: preclinical and clinical studies. Psychopharmacology 164, 121–137 (2002).

144. Huhn, A. S., Berry, M. S. & Dunn, K. E. Review: Sex-Based Differences in Treatment Outcomes for Persons With Opioid Use Disorder. Am. J. Addict. 28, 246–261 (2019).

145. McKee, S. A. & McRae-Clark, A. L. Consideration of sex and gender differences in addiction medication response. Biol. Sex Differ. 13, 34 (2022).

146. Bekhbat, M. & Neigh, G. N. Sex differences in the neuro-immune consequences of stress: Focus on depression and anxiety. Brain Behav. Immun. 67, 1–12 (2018).

147. Barko, K. et al. Brain region- and sex-specific transcriptional profiles of microglia. Front. Psychiatry 13, 945548 (2022).

148. Zannas, A. S., Wiechmann, T., Gassen, N. C. & Binder, E. B. Gene-Stress-Epigenetic Regulation of FKBP5: Clinical and Translational Implications. Neuropsychopharmacology 41, 261–274 (2016).

149. Levran, O. et al. Stress-related genes and heroin addiction: a role for a functional FKBP5 haplotype. Psychoneuroendocrinology 45, 67–76 (2014).

150. Nylander, I. et al. Evidence for a Link Between Fkbp5/FKBP5, Early Life Social Relations and Alcohol Drinking in Young Adult Rats and Humans. Mol. Neurobiol. 54, 6225–6234 (2017).

