## Supplemental Methods and Results for "Single nuclei transcriptomics in human and non-human primate striatum implicates neuronal DNA damage and proinflammatory signaling in opioid use disorder"

###### Human Subjects

Postmortem human brain samples were obtained, following consent from the next of kin, during autopsies conducted by the Allegheny County Office of the Medical Examiner (Pittsburgh, PA). Consent was obtained from next-of-kin and procedures were approved by the University of Pittsburgh’s committee for Oversight of Research and Clinical Training Involving Decedents and Institutional Review Board for Biomedical Research. An independent committee of clinicians made consensus, lifetime DSM-IV diagnoses for each subject using the results of an expanded psychological autopsy, including structured interviews with family members and review of medical records, as well as toxicological and neuropathological reports^151^. The same approach was used to confirm the absence of lifetime psychiatric and neurologic disorders in the unaffected comparison subjects. All procedures were approved by the University of Pittsburgh Committee for Oversight of Research and Clinical Training Involving Decedents and Institutional Review Board for Biomedical Research.Each subject meeting diagnostic criteria for OUD at the time of death (n = 6) was matched with an unaffected comparison subject (n = 6) for sex and as closely as possible for age and PMI (see Table S5). The duration of illness for each OUD subject was at least four years prior to death.

For all subjects, the caudate and putamen were identified on fresh-frozen coronal tissue blocks using anatomical landmarks and tissue was collected via cryostat, using an approach that minimizes contamination from white matter and other striatal subregions and ensures RNA preservation. Fresh-frozen, right hemisphere coronal tissue blocks containing the body of the caudate and putamen, inclusive of plates 24-30 in the rostro-caudal axis, were included for analysis^152^. The rostral face of each block was scored using a #11 scalpel blade while mounted in the cryostat. Sections were cut at 40 µm thickness, and each striatal subregion from an individual section was placed into its respective collection tube until a total of volume of ~50 mm^3^ was collected from each region. This approach minimizes contamination from white matter and other striatal subregions and ensures RNA preservation. The medial-lateral border between the caudate and putamen was scored to exclude the internal capsule. The lateral border of the putamen was defined by the external capsule. The ventral border of the caudate and putamen was defined by the anterior thalamic radiation of the internal capsule or the anterior commissure.

###### Isolation of nuclei from human postmortem brain tissue and library preparation:

Nuclei were isolated from 24 biospecimens of frozen human postmortem brain tissue (12 subjects (Unaffected/OUD) x 2 brain regions = 24 samples). Samples weighing 10-15mg were homogenized using ~10 strokes per glass pestle in 7mL glass douncers with 5mL nuclei isolation medium containing DAPI . Homogenate was filtered using a 40um mesh strainer (FisherScientific #48680). Nuclei were sorted for DAPI fluorescence using a BD FACS Aria at the Boston University Flow Cytometry Core. Approximately 100,000 nuclei were sorted into 7ul of 0.04% bovine serum albumin (MilliporeSigma #126615) in phosphate buffered saline (ThermoFisher #10010031). Nuclei were counted using a hemocytometer and assessed for concentration and debris. 7,000 nuclei were targeted per sample except for one sample with lower concentration where 5,000 nuclei were targeted. The 10x Chromium process was performed and next generation sequencing libraries were prepared using the 10x genomics single cell 3’ gene expression dual index kit.

Libraries were sequenced at the Boston University Single Cell Sequencing Core. The pool of snRNA-seq libraries were sequenced on 7 Next-seq P3 flow cells with an intermediate re-pooling scheme to optimize for 50-80% sequencing saturation, > 8,000 average UMI per cell. Between sequencing runs, we preliminarily aligned the sequencing reads as outlined below to assess quality per sample and estimate the number of viable nuclei and sample complexity. We identified two samples, C-13291 and C-612, to have low QC metrics due to wetting failures, mean UMI per cell <1,000 and estimated # cells >50,000. These samples were excluded from subsequent re-pooling and further analyses (Supplemental Table 1.3 tab STARsolo QC).

###### Nonhuman Primate Subjects

Eight adult rhesus monkeys (*macaca mulatta*) weighing between 6.0 and 13.0 kg served as subjects in the present study. Subjects lived individually in stainless-steel enclosures under a 12-hr light-dark cycle with side and front visual access to other conspecifics. All subjects had continuous access to water and were fed a diet of High Protein Monkey chow (Purina Mills International, Brentwood, MO), fresh fruit, and vegetables. Environmental enrichment (mirrors, toys, foraging boards, music, etc.) was also provided daily. Subject health and well-being were monitored daily by trained technical and veterinary staff. Animal husbandry and research was conducted in accordance with the guidelines provided by the Institute of Laboratory Animal Resources^153^ as adopted and promulgated by the U.S. National Institutes of Health. The facility is licensed by the U.S. Department of Agriculture and all experimental protocols were approved by the Institutional Animal Care and Use Committee at McLean Hospital.

###### Chronic Morphine Dosing in Nonhuman Primate Subjects

Four subjects (3 male, 1 female; mean age: 12 years, range: 9-18 years) received daily morphine treatment for 5 months (see below); four additional subjects, matched for age, weight, and sex, served as experimental controls (3 male, 1 female; mean age: 14 years, range: 11-21 years). Control subjects had a history of nicotine or cocaine exposure but were drug free for ~1 year prior to tissue collection.

Each morphine-treated subject was trained to come to the front of the enclosure for twice daily intramuscular (IM) injections (<0.5 cc) of morphine sulfate (NIDA Drug Supply Program) dissolved in 0.9% saline. Injections were administered at 09:30 and 17:00 hours. A gradual dosing escalation method (0.5 log unit increase every 3 days) was used to achieve the terminal dosage of 9 mg/kg/day (i.e., 4.5 mg/kg, BID). This dosage was selected to produce moderate opioid dependence^154,155^. All subjects received approximately 1,500 mg of morphine over the 5-6-month period of chronic dosing. On the last day and approximately 3 hrs following the last IM morphine injection, each subject received an IM injection of ketamine (10 mg/kg) followed by 5.0 ml IV of a pentobarbital-based euthanasia solution (Beuthanasia-D).

###### Rhesus brain tissue preparation

After sacrifice, brains were rapidly dissected into slabs and frozen on metal plates in liquid nitrogen vapor. Time between animal sacrifice and tissue freezing was between 1-2 hours for all animals. Brains were kept stored at -80 C until punching, when they were punched on a microtome with guidance from a macaque anatomist (Dr. S. Haber). Punches were stored at -80 C until the day of nuclear isolation and encapsulation.

Single nucleus RNA sequencing in rhesus brain tissue

Nuclei were isolated as in previous studies^156^, with minor modifications. Punches were placed into buffer HB (0.25 M sucrose, 25 mM KCl, 5 mM MgCl_2_, 20 mM Tricine-KOH pH 7.8, 1 mM DTT, 0.15 mM spermine, 0.5mM spermidine, protease inhibitors) and placed in a Dounce homogenizer for 10 strokes each with loose and tight pestles. A 5% IGEPAL solution was added to a final concentration of 0.3% followed by five additional dounce strokes, then the lysate was filtered through a 40-μm strainer. Nuclei were mixed with an equal volume of 50% iodixanol and then layered on top of an iodixanol gradient of 40% and 30% layers in a 2 mL dolphin microcentrifuge tube. Nuclei were spun by centrifuging at 10,000 x g for 4 min at 4°C and then collected by aspiration at the interface of the 30% and 40% iodixanol layers. Nuclear concentration and prep quality were ascertained by loading on a hemocytometer and were diluted to a concentration of 80-100K and 15% iodixanol with Buffer HB prior to loading on InDrops v3 platform. Single-nuclei suspensions were encapsulated into droplets, lysed, and the RNA within each droplet was reverse-transcribed using a unique nucleotide barcode as described previously^157^. Approximately 6,000 nuclei, in two batches of 3,000 nuclei each were processed per library and sequenced on Illumina NovaSeq S2 chips (at a density of approximately 20,000 reads/nucleus).

###### Single nuclei RNAseq data processing

We aligned single nuclei RNA-seq (snRNA-seq) reads to the human genome (GRCh38.p13) or rhesus macaque genome (rheMac10) for each output with the turn-key single-cell transcriptomics method STARsolo, which is folds faster than the CellRanger pipeline and equally accurate (v2.7.9a) ^158^. For the macaque samples, we used a set of gene annotations by mapping the human gene annotations to the rheMac10 genome using the liftoff tool^159^. These alternate rheMac10 gene annotations are deposited to Carnegie Mellon University’s Kilthub repository resource at ref^160^.We chose parameters for the STARsolo UMI quantification to closely replicate the 10X Cell-Ranger pipeline v6 and use the filtered genome and gene annotation available from 10X Genomics (<https://support.10xgenomics.com/single-cell-gene-expression/software/downloads/latest>, Human reference 2020-A). We ran STARsolo to allow for pre-mRNA gene counts as well as exonic counts for nuclear RNA and to separately count introns and exons for RNA velocity analyses, (--soloFeatures GeneFull Velocyto). We used the following parameters to correct cell barcodes, de-duplicate transcripts by their unique molecular identifier (UMI), assign UMI counts to genes, and pre-filter cells that are likely empty droplets (--soloType Droplet --soloCBmatchWLtype 1MM --soloCellFilter EmptyDrops_CR --soloMultiMappers EM --soloUMIdedup 1MM_CR). We pre-process the UMI gene x cell count matrix to reduce inherent biases in the technology. We identified likely ambient RNA contamination with SoupX ^161^, empty droplets with DropletQC ^162^, doublets with scds ^163^, and damaged nuclei with miQC ^164^. For each of these analyses, each sample (GEM well) was analyzed separate from each other. We ran SoupX as described to estimate the fraction of ambient RNA from both raw and unfiltered UMI count matrices from STARsolo and perform ambient RNA removal aware of the cell clusters in the filtered matrix. For just the SoupX analyses, we clustered the cells with Seurat v4 ^165^ with FindClusters(algorithm = 2, resolution = 0.5). For DropletQC, we used the intronic and exonic UMI counts per cell per gene from STARsolo to get the fraction of intronic UMI per cell (referred to as the nuclear fraction). We identified empty droplets with default DropletQC parameters (nf_rescue = 0.50, umi_rescue = 1000). We identified droplets with scds’s hybrid algorithm using the function cxds_bcds_hybrid to estimate doublet scores and called doublets on cells with scds.hybrid_score > 1.0. We identified damaged cells with high percentage of mitochondrial UMI counts using only miQC which uses a Bayesian EM algorithm to learn the relationship between mitochondrial UMI counts and number of captured genes. We used the posterior probability cutoff of 0.75 to call damaged cells by miQC.

To combine cells together across samples, we normalize the UMI counts with the variance-stabilizing SCTransform and glmGamPoi on each sample ^166,167^ and jointly embed cells across samples with reciprocal PCA integration ^165^ as outlined in <https://satijalab.org/seurat/articles/integration_rpca.html>. In this joint embedding, we over-clustered the dataset with FindClusters(algorithm = 2, resolution = 1)and removed any cluster with more than 10% of cells flagged by miQC, scds, or DropletQC as low-quality biased-clusters in the data.

###### Labeling striatum cells with a macaque snRNA reference dataset

We annotated our cells from the human or monkey striatum to a recently published high-resolution snRNA-seq reference dataset of the non-human primate striatum ^37^ using Seurat v4. We downloaded the monkey snRNA-seq processed, annotated gene UMI counts for all cells and MSNs from GSE167920. *He, Kleyman et al.* had aligned the snRNA-seq reads to the rheMac10 genome using the GRCh38 gene annotation liftover to rheMac10, so the gene-wise labels represent the UMI counts on the rheMac10 genome most orthologous to human. For both full nuclei and MSN subset datasets, we re-processed the macaque cells with SCTransform , glmGamPoi, and reciprocal PCA with default parameters as above to enable label transfer using the most recent integration algorithms in Seurat.

To transfer cell annotations from the reference macaque striatum dataset to the human or macaque striatum cells, we perform two label transfers at increasing resolutions: one with all cells and another with just MSNs. As *He, Kleyman et al.* described, the differences between transcriptionally and anatomically distinct MSN subtypes are subtle, so we split the annotations into two steps to optimally annotate the cells. The first label transfers the cell classes (Oligodendrocytes, MSNs, Interneurons, etc.) from the macaque to the human dataset with the Seurat functions FindTransferAnchors(reduction = ‘rpca’) and TransferData. Next, we identified cells or cell clusters that were labeled as MSNs and transferred MSN subtype labels (D1.Striosome, D2.Striomsome, etc.) from the macaque to human datasets. We filtered out cells where the cell class or cell subtype labels have max prediction scores less than 0.5 as these tend to represent noisy predictions due to low quality cells from either datasets. We confirmed accurate label transfer at the cell class and cell subtype levels with published marker genes and similar proportions across subjects and samples.

Even with the robust cutoffs that we applied to this dataset to remove likely low quality or doublet cells, we find a residual subset of the data that contain these cells. Upon clustering, doublet cells tend to project into the UMAP space as long streaks between two well-defined cell types. Low-quality cell types would project into the UMAP space as amorphous cell types without clear boundaries. Using these embedding features, we selected these clusters with Seurat’s FindClusters(resolution = 1) function, confirmed that they have the indicative QC metrics, and removed them from analyses.

###### Annotating interneurons with mouse marker genes

To demonstrate the high resolution of our datasets, we annotated the striatal interneuron using previously characterized mouse markers of these subtypes ^133^. We sub-clustered the interneurons labeled by the macaque dataset and annotated them manually as interneuron subtypes best labeled by the marker genes *TH*, *PTHLH*, *SST*, or *CCK*. The previously published macaque snRNA-seq dataset had too few of these interneurons sampled from N=2 monkeys, so these subtypes were under-represented to be sub-clustered. In this study, we sampled more broadly from N=12 individuals (human) or N=8 individuals (rhesus macaque). While these interneuron subtypes are clearly distinct in this dataset and in the neural circuits, they still are under-represented to power certain downstream. For these analyses, we analyze these cells together as “Interneurons”.

###### Differential gene and cellular state expression analysis in humans

To investigate the gene expression differences in OUD and unaffected individuals, we used the pseudo-bulk aggregation of gene expression profiles. Many have shown that pseudo-bulk based case-control differential expression analyses robustly detect gene-level differences with lower false discovery due to repeated measures from single cells of the same individual^168–170^. The raw UMI counts were added together from the same individual, brain region, and cell type to create the pseudo bulk profiles. We aggregate the interneuron subtypes together as “Interneurons”. We filtered out pseudo-bulk profiles aggregated from more than 20 cells. We filtered out mitochondrial, ribosomal, and low-expressing genes with less than 5 average UMI counts. We retained 20,203 genes and 210 pseudobulk profiles to apply the voom-limma method^171^ for differential gene expression analyses and the sva method to construct surrogate variables to identify un-modeled sources of transcriptomic variation. These statistical methods together address several challenges in analyzing cell type differential expression across multiple axes of meaningful biological variation: 1) accounting for correlated pseudo-bulk samples shared across individuals with the duplicateCorrelation(block = Subject.ID), 2) estimating quality weights for adjusting for cell type proportions with the function voomWithQualityWeights(), and 3) calculating un-modeled variation in high-dimensional single cell data with the sva()function. To plot the gene expression profiles, we use the normalized counts per million (CPM) of each pseudobulk profile corrected for the batch effects and surrogate variables unrelated to the OUD diagnosis using the cleaningY() function (<https://github.com/LieberInstitute/jaffelab>). For heatmap visualizations of the gene expression profiles, we also z-normalized the corrected expression profile of each gene grouped by cell type, since gene expression is highly cell-type specific.

We calculated the differential expression in OUD vs. unaffected comparison individuals for two sets of hypotheses:

1) What is the differential expression in each cell type averaged across brain regions and sex?

2) What is the differential expression within the subset of female or male individuals?

3) What is the differential expression interaction between sex and OUD?

To achieve these comparisons, we used one linear model with a nested variable capturing the interaction between cell type, OUD diagnosis, Sex, and Region, Celltype_Dx_Sex_Region. The nested variable allows for contrasting subsets of the data to calculate differential expression for each family of hypotheses at different cell type resolutions: ~ 0 + Celltype_Dx_Sex_Region + Age + PMI + RIN + GDR + 23 SVs, (GDR: gene detection rate, SV: surrogate variables). We estimate the effect of OUD vs. unaffected individuals across each annotated cell type, across neuronal subtypes (Neuron), across glial types (Glia), and across all cell types (All). For the first set of hypotheses at the annotated cell type level, we used the simple contrast CelltypeA_OUD_Sex?_Region? - CelltypeA_UC_Sex?_Region? with samples from both Sex and Regions to obtain the average effect across those variables (?, the wildcard placeholder for F, M, Caudate, or Putamen). For the second set hypotheses, we only included individuals within each Sex to obtain the effect of OUD vs. unaffected within each sex subset. For the third set of hypotheses, we use the following contrast: (CelltypeA_OUD_SexF_Region? - CelltypeA_UC_SexF_Region?) - (CelltypeA_OUD_SexM_Region? - CelltypeA_UC_SexM_Region?).

We calculated the differential expression in chronic morphine exposure vs. non-morphine treat rhesus macaque subjects. We used one linear model with a nested variable capturing the interaction between cell type, morphine treatment and matching stats. ~ 0 + Celltype_Tx + Pair + SVs. We estimate the effect of morphine treatment vs. no-morphine individuals across each annotated cell type, across neuronal subtypes (Neuron), across glial types (Glia), and across all cell types (All). For the first set of hypotheses at the annotated cell type level, we used the simple contrast CelltypeA_morphine - CelltypeA_control.

###### Gene set enrichment analyses

We identified pathways that are differentially altered in OUD using gene set enrichment analyses with the molecular signatures database ^172^ and the fgsea and msigdbr R-packages ^173^. We included the hallmark gene pathways, curated gene sets from BioCarta, KEGG, Canonical Pathways, Reactome, and WikiPathways, and ontology gene sets. We also downloaded and included curated pathways from SynGO in enrichment analyses^174^. We report the full list of enriched pathways corrected for multiple hypotheses within each set of hypotheses in Table S7. We clustered redundant/related pathways using custom R-scripts and the igraph R-package creating networks of pathways connected by overlapping genes as previously described ^175–177^. We visualize the clustered pathways using igraph functions and report both clustered and singleton pathways alongside each figure in tables S7, S9, and S11. For the interaction score for calculating pathway-enrichments between female- and male-biased pathway analyses, we use the log2(fold-change) and the p-values from the OUD v. UC effect within females or males differential analyses to calculate an interaction score: sign(log_2_FC_F_) * (-log_10_(p-value_F_)) - sign(log_2_FC_M_) *( -log_10_(p-value_M_)). We overlap enriched pathways using this calculation vs. the pathways enriched in the standard OUD v. UC in females or males to further support the interpretation of female- or male- biased pathways.

To intersect the genes found to be differentially expressed in OUD within females and males, we also compared to differentially expressed genes identified by microarray studies of striatum of male mice exposed to drugs^124^. Piechota et al. identified from hierarchical clustering 2 main gene signatures that became pronounced time lapsed since drug exposure, A and B, with subsets of the second B1, B2, and B3. We identified the human orthologs of these gene sets and performed GSEA as above on the differentially expressed genes identified across the full cohort, on the subset of female and male subjects, or the caudate and putamen samples. We report the main findings in the text and the full set of enrichments in Table S16.

###### Transcription factor-gene regulatory network analyses

The single nuclei RNA-seq dataset in this study provides a new resource to generate gene pathways. To discover new pathways from our single cell RNA-seq, we applied the pySCENIC Protocol to build transcription factor (TF)- gene regulatory networks ^49,178^. To potentially capture individual-specific TF-gene regulatory relationships, we ran the GRNBoost2 with multiprocessing to infer TF-gene relationships from the pySCENIC package on each individual separately ^178,179^. To speed up the GRNBoost2 run time, and reduce bias by cell type proportion, we downsampled oligodendrocytes (~60% of all cells) to the next most prevalent cell type, Astrocytes (~8%), reducing the GRNBoost2 run time by 50%. Furthermore, we subset to the ~20k genes that are expressed and analyzed in the pseudobulk differential expression analysis further reducing GRNBoost2 runtime by 30%. Since GRNBoost2 is a stochastic method, we ran this step 3-4 times for each individual and selected the TF-gene relationships that were reproducible >80% across runs. Similarly, to aggregate the TF-gene relationships across individuals, we aggregated TF-gene relationships that were detected in 80% of 12 individuals. Across these aggregation steps, we averaged the importance scores for selected TF-gene relationships. We selected likely direct TF-gene relationships by filtering relationships where the gene’s promoter contain the TF’s binding motifs using the cisTarget method implement in pySCENIC package using the pre-built TF-motif promoter binding databases for human accessed from <https://resources.aertslab.org/cistarget/databases/> in January 2023^49^. We report the list of discovered TF-gene relationships in the table S17.

###### Analyses of cellular activation of gene sets

We calculated the level of activity of each collection of gene sets with the AUCell method as described previously^49^ on the list of TF-gene regulatory networks or custom gene sets (described later). For the TF-gene relationships identified above, we calculated the differentially active TF-gene modules between unaffected and OUD individuals by creating the pseudobulk average of each TF-module by individual, brain region, and cell type. We performed linear regressions to assess the effect of OUD diagnosis accounting for covariates RIN, Age, Sex, Region, and number of aggregated cells. We corrected multiple testing of the OUD effect on TF-gene module activity across all cell types and TF modules and reported the full cell type by sample linear regression are reported in Table S8. We similarly performed a similar pseudobulk average at the biological sample level aggregating over individual and brain regions. We visualized the significant TF-gene module activity differences set at FDR < 0.05 in glia and neurons and of select modules with the igraph R-package^177^.

In addition to the TF-gene relationships, we assessed the state of DNA damage in OUD using two published gene signatures of DNA damage^13^. The neuronal DNA damage signatures were collected by Welch et al. from fluorescence activated sorting of mouse brain nuclei gated for NeuN+:gH2AX+ nuclei. Using the human orthologs of these mouse genes, we scored our single nuclei RNA-seq datasets with the AUCell method to study the differences in DNA damage gene signatures between unaffected and OUD individuals or no-morphine or chronic morphine treated rhesus macaque individuals. We applied the same pseudobulk average of DNA damage scores and reported the linear regression interaction between OUD and cell type in DNA damage signatures in Table S10. We similarly apply the same pseudobulk average of DNA damage scores and report the linear regression interaction between chronic morphine exposure and cell type regressing out the match pair in DNA damage signatures.

hdWGCNA

hdWGCNA was used to further analyze gene co-expression networks across striatal cell types. Each of 6 cell types of interest (D1-Striosome, D2-Striosome, D1-Matrix, D2-Matrix, D1-D2 hybrid and microglia) was used as input. Gene expression data was SCT transformed. Each dataset was then collapsed into a single metacell. Softpower was selected for each of the metacell to construct gene co-expression networks. The gene modules were identified using unsupervised clustering via the Dynamic TreeCut algorithm with default settings. For each module, Top 100 hub genes for each module were identified. Eigengene of identified modules was correlated with “traits” (in our case, OUD, Sex, Race and RIN, number of features, number of RNA count and percent of mitochondrial RNA) using Pearson correlation. Modules with significant correlation (p<0.05) with OUD will be used for downstream analysis, especially, modules significant with OUD (Figure S9,10,11; Table S18,S19) were prioritized. For cell types with no module uniquely correlated with OUD, hub genes from all OUD trait correlated modules were used as input for pathway analysis. Metascape (metascape.org) was used for pathway analysis on modules from co-expression gene networks by each cell type (Figure S11; Table S18).

Statistical Analyses

We correct for multiple hypotheses tested in this study using the false discovery rate (FDR) less than alpha = 0.05. In different expression analyses, we perform FDR correction across all cell types tested. Similarly for gene set enrichment analysis and AUCell pseudobulk analyses, we perform FDR correction across all pathways and differentially expressed genes across cell types. We compute the FDR correction with the swfdr R package which increases power by leveraging Storey’s q-value and modeling the relationship of null P-values and independent variables such as the average logCPM or the number of genes in a pathway^180,181^. In differential gene expression analyses, we calculate the FDR within each cell type comparison (adj.P.Val.Within) and between all cell type comparisons (adj.P.Val.Between). We report the more conservative correction between cell types in the main text and figures, adj.P.Val.Between. Measured metrics (gene expression counts, quality control metrics) or derived metrics (AUCell gene activity scores) are repeated measurements of the same cohort of human or rhesus macaque individuals. Metrics that are differentially expressed across multiple cell type conditions are reported and summarized at the largest significant adjusted P-value and lowest effect size magnitude. Genes or metrics that are only significantly differentially expressed across one cell type are reported with exact adjusted P-values and effect sizes rounded to 2 significant digits. Exact P-values and effect sizes are reported in the supplementary data. All statistical analyses performed use two-sided test statistics.

For human samples, linear regressions of differential AUCell modules, or other quality control metrics at the pseudobulk analyses were performed across biospecimens (N=22 biospecimens) controlling for relevant covariates including brain region extracting the interaction between OUD diagnosis with cell type. The general linear model for this linear regression for sample level is ~ Dx + Age + PMI + RIN + Sex and for cell type level is ~ Dx:Celltype + Celltype + Age + PMI + RIN where we extract the interaction term to determine OUD diagnosis-specific changes in a cell type. For rhesus macaque samples, linear regressions of differential expression and differential AUCell modules were performed across biospecimens (N=8 biospecimens) controlling for matched pair grouping variable, which accounts for sex, age, and weight. The general model for this linear regression at the individual level is ~ Tx + Pair and for cell type level is ~ Celltype:Tx + Celltype + Pair.

##### Supplementary Tables

###### Table S1. Comparison of quality control metrics by OUD diagnosis

False discovery rate-corrected t-tests comparing aggregated single nuclei RNA-se quality control metrics between UC and OUD individuals across the caudate or putamen.

###### Table S2. Comparison of proportion of cell types by OUD diagnosis

False discovery rate-corrected multiple mixed effect linear regression for any difference in cell type proportion between UC vs. OUD controlling for covariates as fixed effects and individual as a random effect.

###### Table S3. Significant marker genes of striatal cell types

Significant marker genes for each striatal cell type using the Seurat FindMarkers() function with default parameters to compute one-vs-all comparison using Wilcoxon-rank sum test. We computed the FDR across all cell types and significant marker genes are reported at FDR < 0.05.

###### Table S4. Significant marker TF-gene modules of striatal cell types

Significant marker TF-gene module activation for each striatal cell type using t-tests comparing the TF-gene module AUCell scores of each cell type with all others. The false discovery rate was computed across all cell types and significant marker genes are reported at FDR < 0.05.

###### Table S5. Postmortem individual-level and tissue-level metadata

Description of the biospecimen-level and individual-level metadata describing relevant covariates and opioid use disorder diagnoses.

###### Table S6. Differentially expressed genes in UC vs. OUD within striatal cells

Outputs of differential expression across striatal cells or the aggregate “meta cells”, All, Neuron, and Glia. The differential expression FDR was calculated using the swfdr method to increase power by modeling power with an independent factor, average expression. We calculated FDR within each cell type comparison (adj.P.Val.Within) and between all cell type comparisons (adj.P.Val.Between) and report the more conservative correction between cell types in the main text and figures, adj.P.Val.Between.

###### Table S7. Enriched pathways in UC vs. OUD, main comparison

Outputs of the gene-set pathway enrichment analyses using the main differential expression from UC vs. OUD. We calculated FDR across all cell type comparisons using swfdr modeling power by the number of genes in a gene set.

###### Table S8. Enriched TF-gene modules in UC vs. OUD

Outputs of differential pseudo-bulk average transcription factor-gene regulatory network activation in striatal cell types using linear regression to control for covariates.

###### Table S9. Clustered neuronal pathways in UC vs. OUD, main comparison

Neuronal pathways from Table S7 with additional clustering to identify related, enrich pathways. The cluster number corresponds to those in Figure 2B-C.

###### Table S10. Comparison of neuronal DNA damage score by OUD diagnosis

Outputs of differential pseudo-bulk average DNA damage score activation in striatal neurons using linear regressions to control for covariates.

###### Table S11. Clustered glial pathways in UC vs. OUD, main comparison

Glial pathways from Table S7 with additional clustering to identify related, enrich pathways. The cluster number corresponds to those in Figure 4B-C.

###### Table S12. Differentially expressed genes in UC vs. OUD subset to female or male

Similar outputs to Table S6 from differential expression across striatal cells or the aggregate “meta cells”, All, Neuron, and Glia. These differential expression tables assess UC vs. OUD in female or male individuals (SexF and SexM, respectively).

###### Table S13. Enriched female-biased pathways in UC vs. OUD

Similar outputs to Table S7 showing gene-set pathway enrichment analyses using the differential expression from UC vs. OUD that were biased to female individuals with OUD.

###### Table S14. Enriched male-biased pathways in UC vs. OUD

Similar outputs to Table S7 showing gene-set pathway enrichment analyses using the differential expression from UC vs. OUD that were biased to male individuals with OUD.

###### Table S15. Enriched pathways with a sex interaction in UC vs. OUD

Similar outputs to Table S7 showing gene-set pathway enrichment analyses using the differential expression from UC vs. OUD that had a differential interaction score between DEGs in females vs. in male individuals.

###### Table S16. Enriched pathways from Piechota et al. 2010

Gene-set enrichment analysis using gene sets from Piechota et al. 2010, gene pathways after drug exposure in male mouse striatum, with differentially expressed genes.

###### Table S17. Detected transcription factor (TF)-gene relationships

The transcription factor-gene regulatory relationships identified using the pySCENIC protocol to build gene regulatory networks. These TF-gene relationships were identified across multiple runs and across multiple individuals with and without OUD.

###### Table S18. Significantly enriched pathways from gene co-expression modules significantly correlated with OUD in medium spiny neuron subpopulations and microglia.

###### Table S19. Weighted gene co-expression analysis (WGCNA) across medium spiny neuron subpopulations and microglia.

##### Supplementary Figures

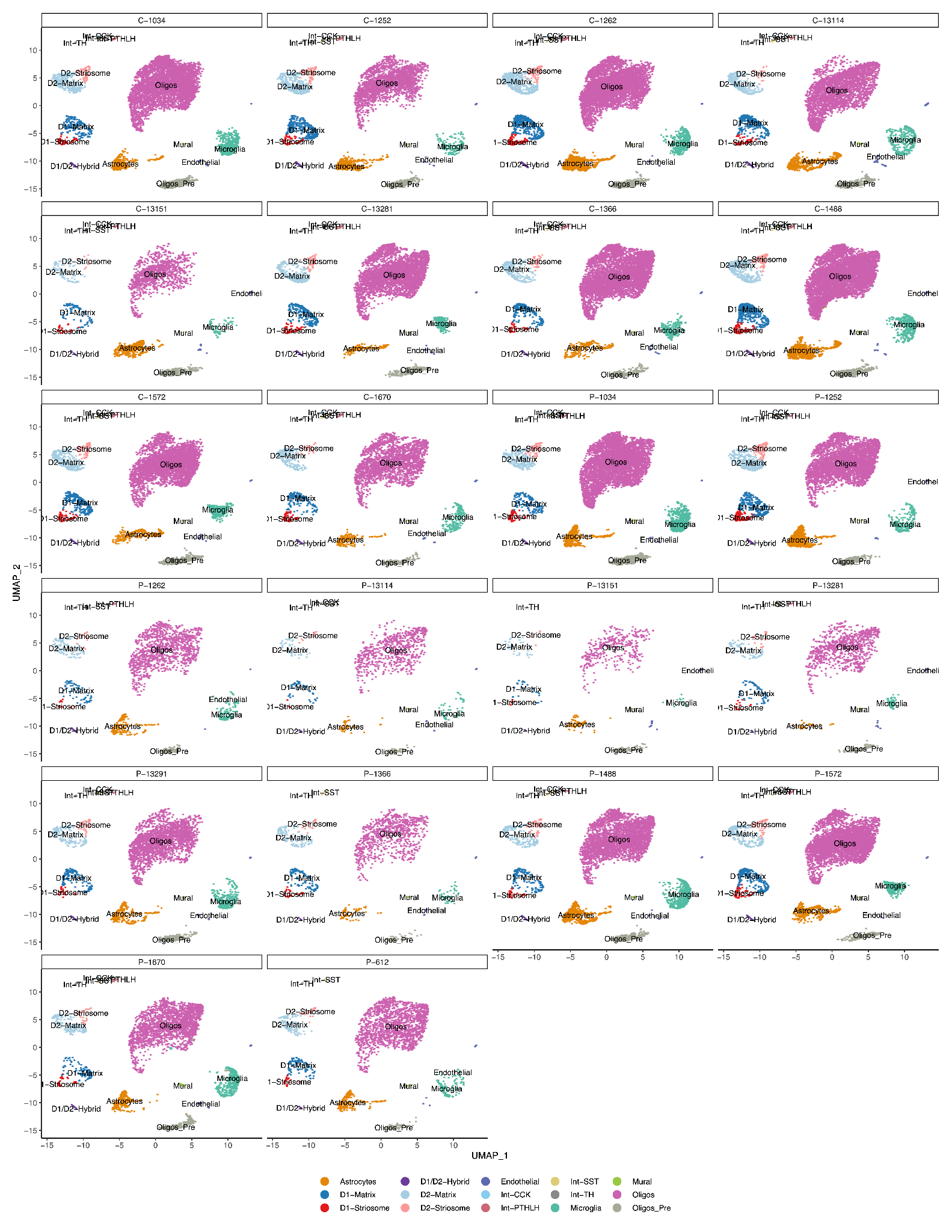

###### Figure S1. Low dimensionality projection of striatal cell types after QC filtering and annotation.

###### Cell clusters are based on marker genes, with each cell cluster represented by a different color. Cell clusters are plotted for each subject (number) and tissue type (C: caudate; P: putamen).

###### Figure S2. Marker genes for dopamine receptor subtype medium spiny neuron subpopulations.

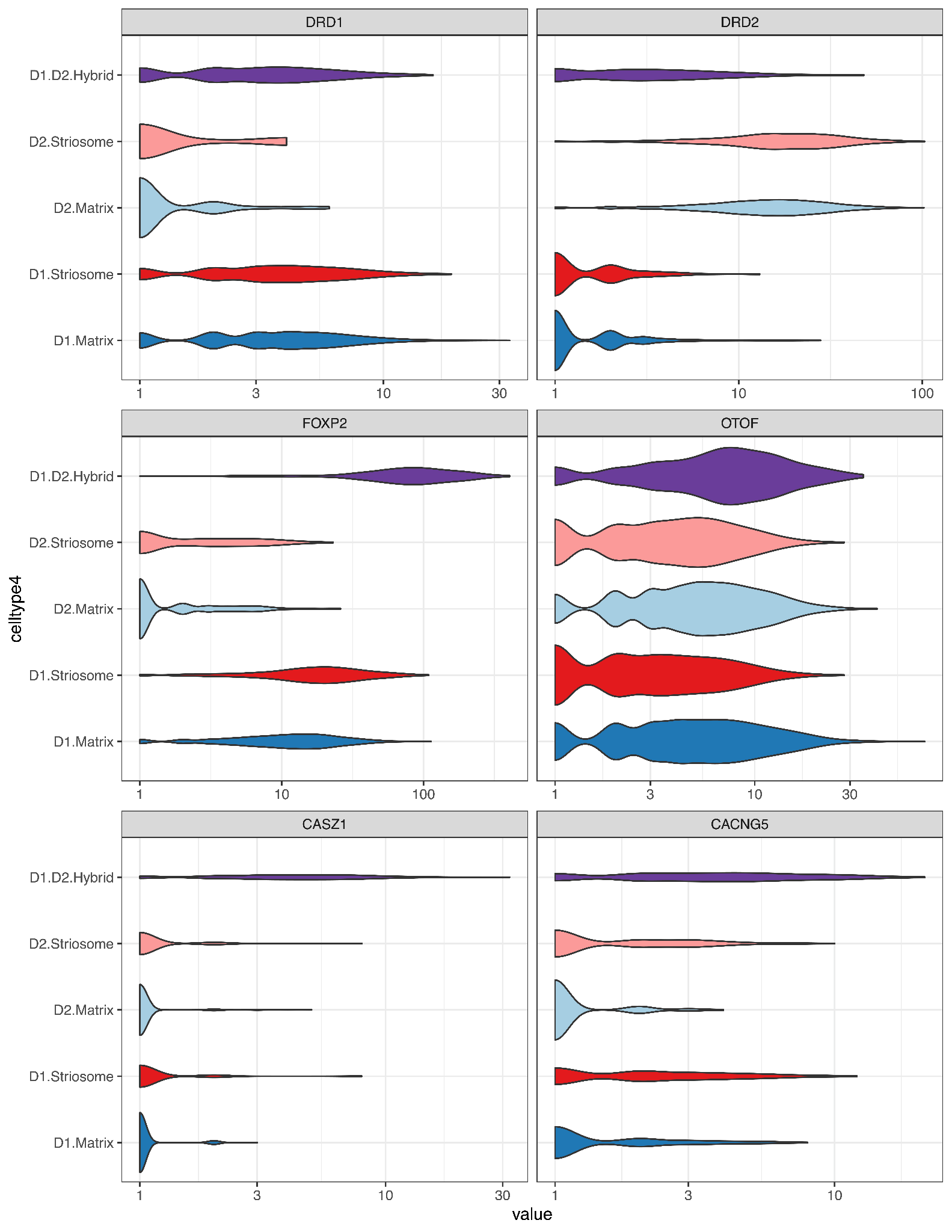

###### Violin plots showing normalized counts for marker genes by major dopamine receptor medium spiny neuron subtype. The mouse orthologs of OTOF, CASZ1, and CACNG5 are marker genes for a previously described striatal cell type differently termed “eccentric spiny projection neuron”, D1-hybrid, or D1-*Pcdh8* neurons.

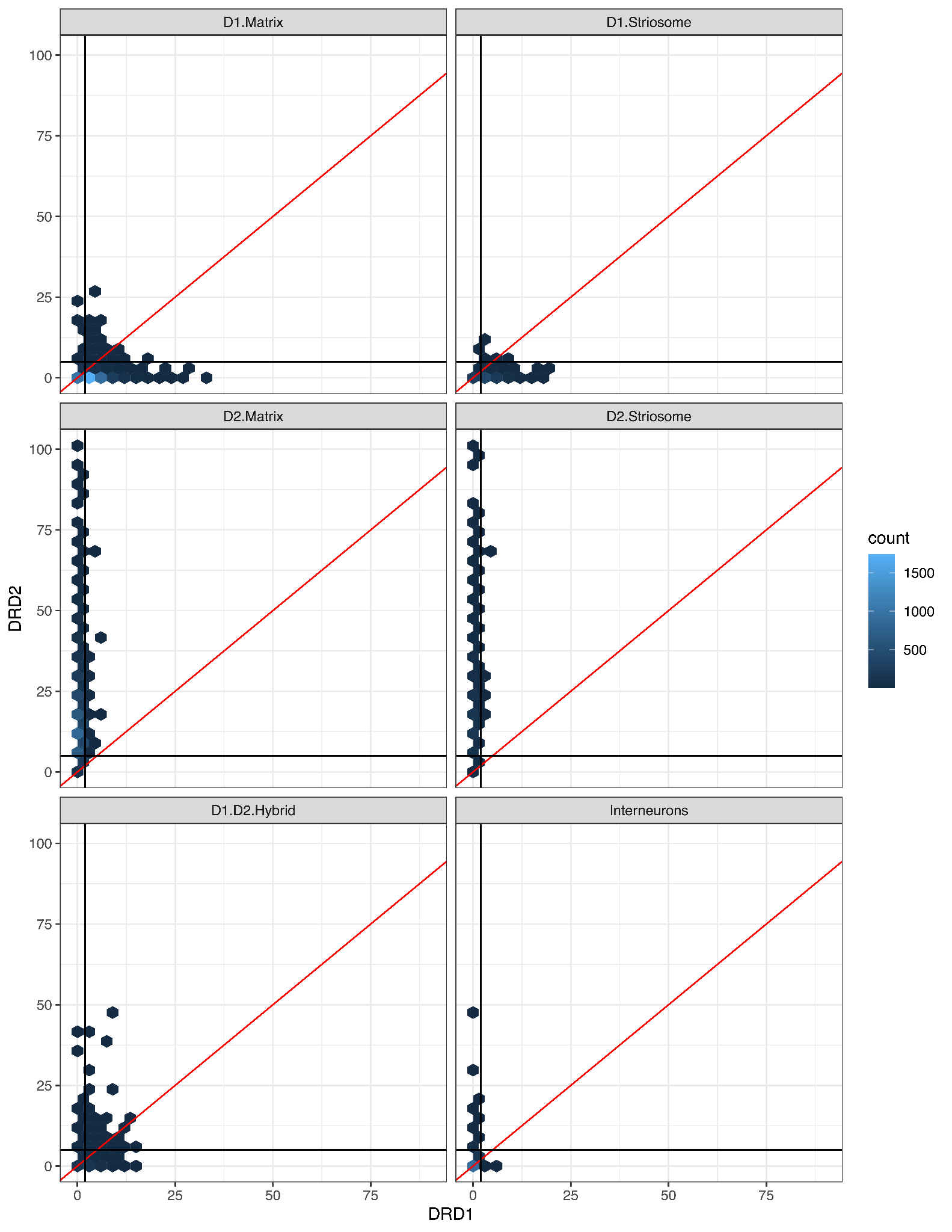

###### Figure S3. Co-expression heatmap of single cells expressing *DRD1* and *DRD2* in major neuronal subpopulations.

###### Gene expression counts for *DRD1* on x-axis and *DRD2* on y-axis. Co-expression profiles indicates comparatively more consistent expression of both *DRD1* and *DRD2* in D1/D2-hybrid medium spiny neurons.

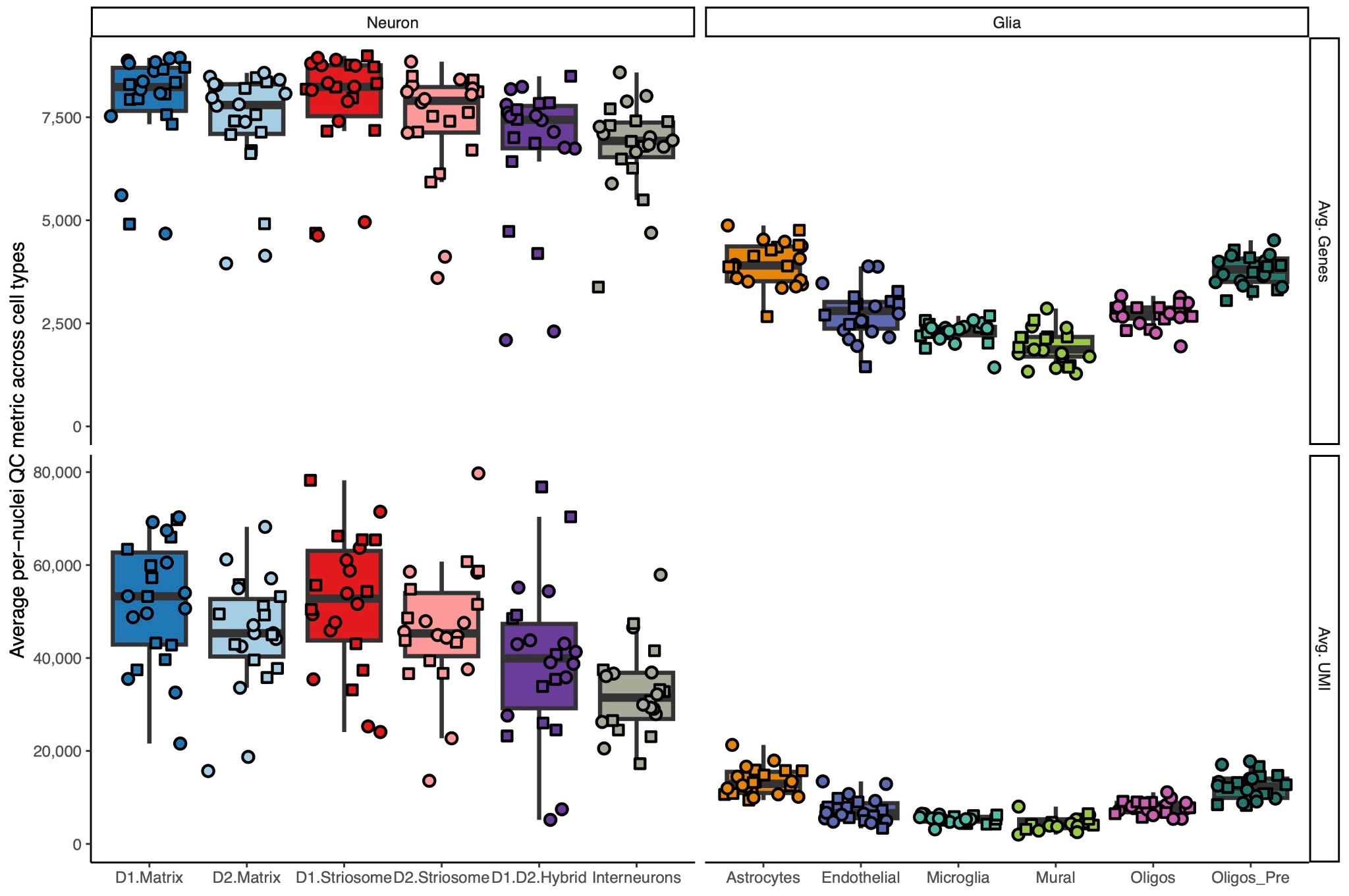

###### Figure S4. Cluster-specific quality control metrics of striatal cell types after QC filtering and annotation.

###### Quality control metrics (average number genes per cell and UMIs per cell) show neuron and glia-specific differences in sample quality. Each point is a unique biospecimen. Round points represent female and squares represent male subjects.

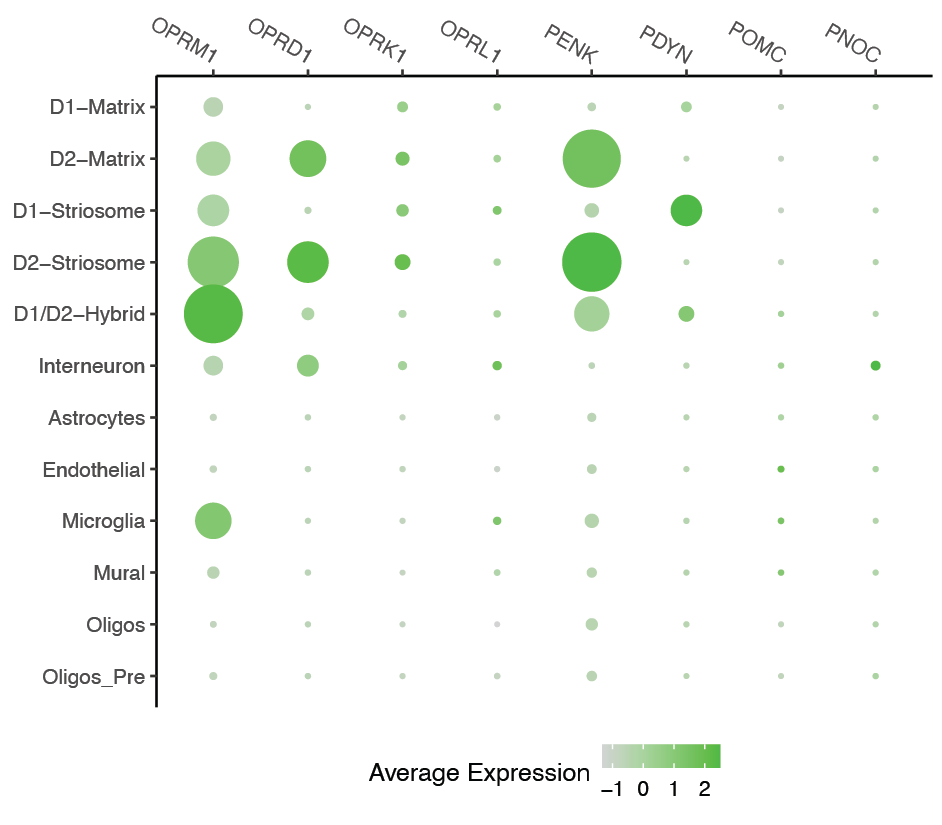

###### Figure S5. Expression patterns of striatal opioid receptor and endogenous ligands

Dot plots of the opioid receptor and endogenous ligands across the annotated striatal cell types. The normalized expression patterns are averaged across all cells and subjects.

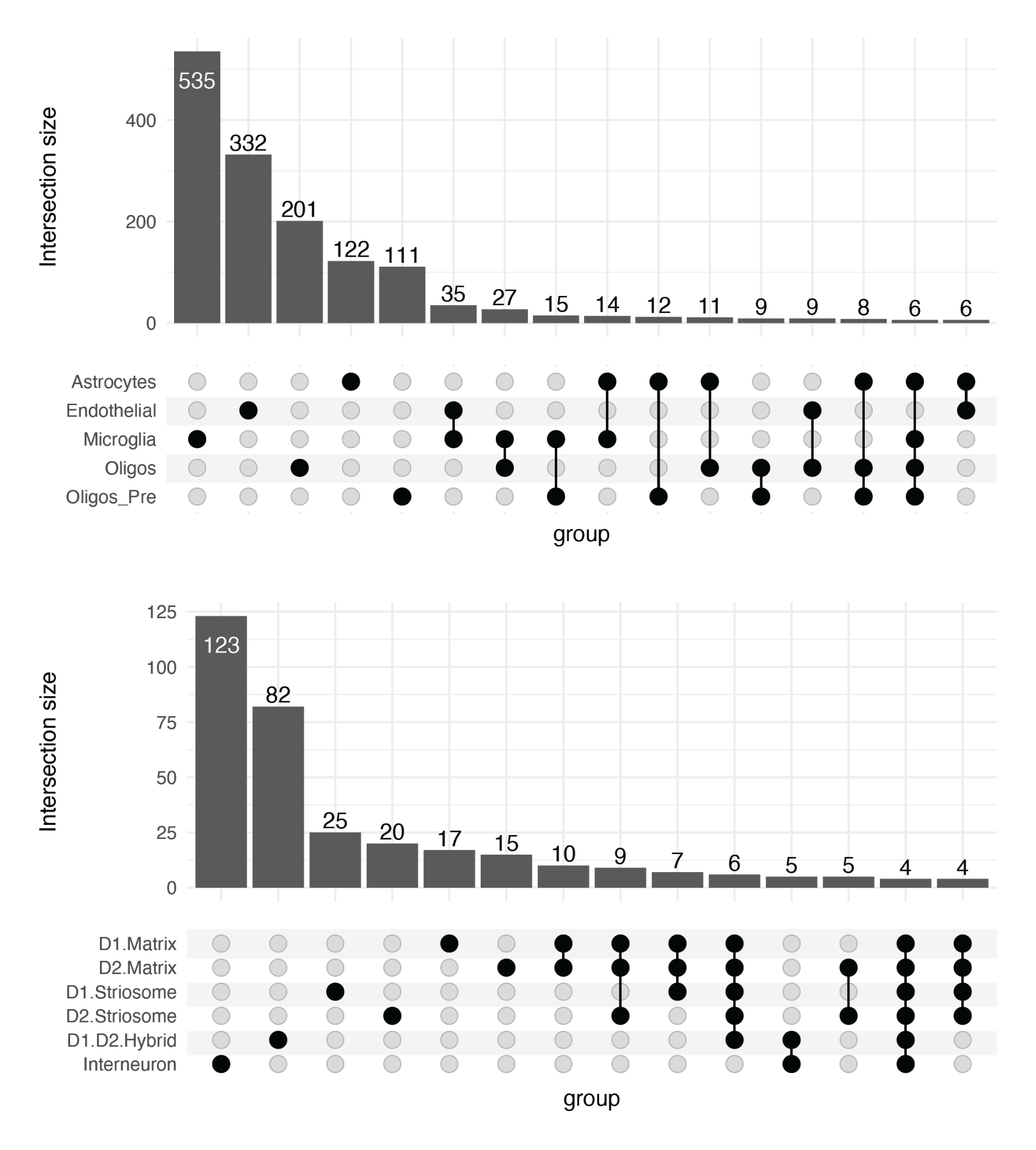

###### Figure S6. Upset plot of overlapping differentially expressed genes in OUD

Upset plot showing the overlaps in significantly differentially expressed genes (FDR < 0.05) across glia (top) or neurons (bottom). The histogram shows how many genes are in each intersection of cell types that share DEGs.

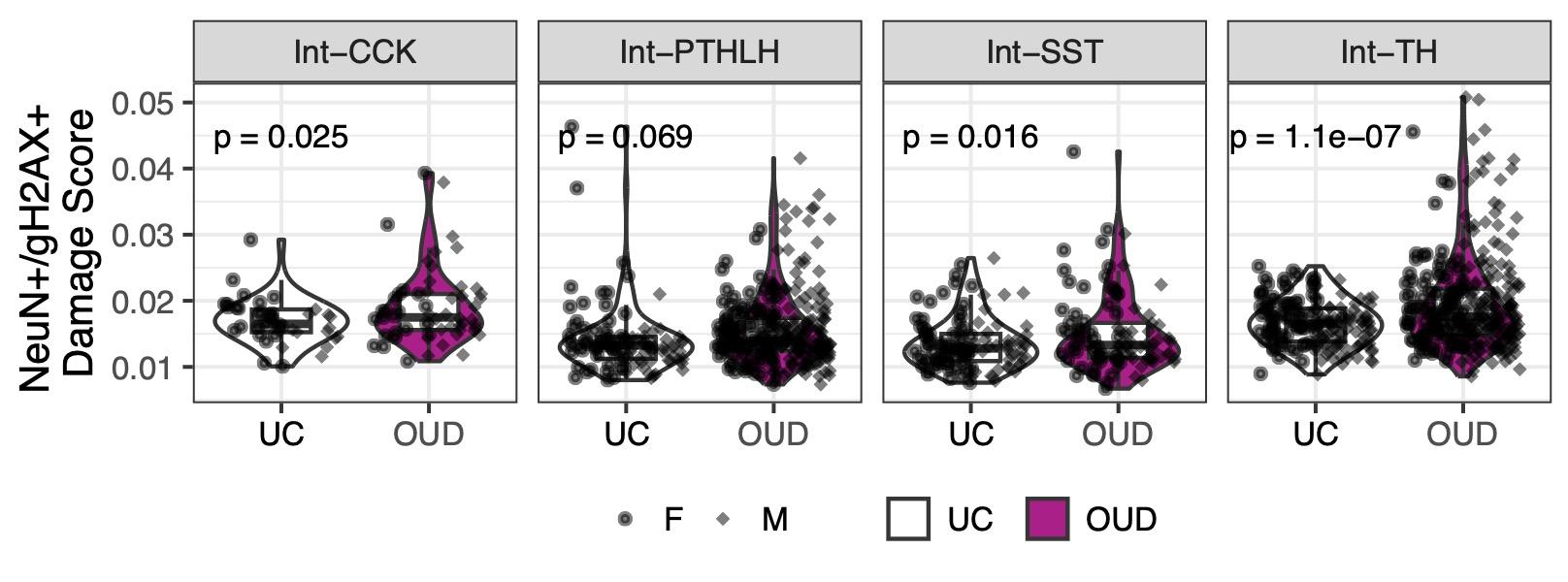

Figure S7. Elevated DNA damage markers across interneuron subtypes associated with OUD. Boxplot of subject-level average DNA damage scores across striatal interneuron subtypes between unaffected and OUD subjects (linear regression comparing unaffected, UC, to opioid use disorder (OUD) subjects).

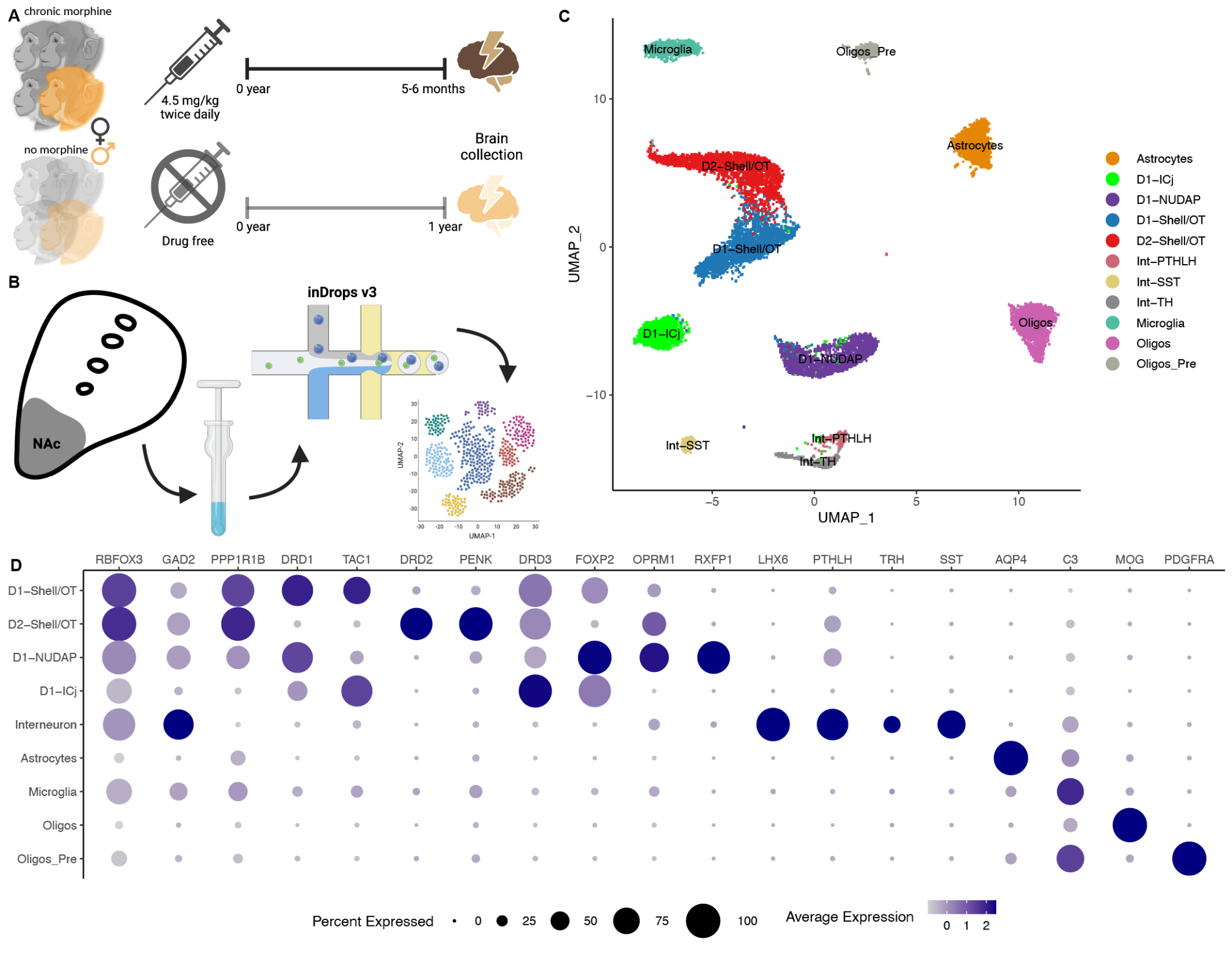

###### Figure S8. Chronic morphine exposure and single nucleus RNA-seq of the rhesus macaque striatum

(A) Schematic of chronic morphine dosing strategy for 5-6 months prior to brain tissue collection in N=4 rhesus macaques to a total of 1,500 mg of total morphine exposure. N=4 control treatment subjects are matched by age, sex, and weight to the chronic morphine cohort and are unexposed to morphine or other drugs for 1 year prior to brain tissue collection. (B) Schematic of nucleus extraction from a tissue punch of the nucleus accumbens for inDrops single nuclei RNA-seq library synthesis and clustering. (C) UMAP projection of post-quality control and annotation of the rhesus macaque striatal cell types using a high quality reference. (D) Dot plot showing the marker genes used to validate the annotations of the chronic morphine rhesus macaque striatal single nucleus RNA-seq dataset.

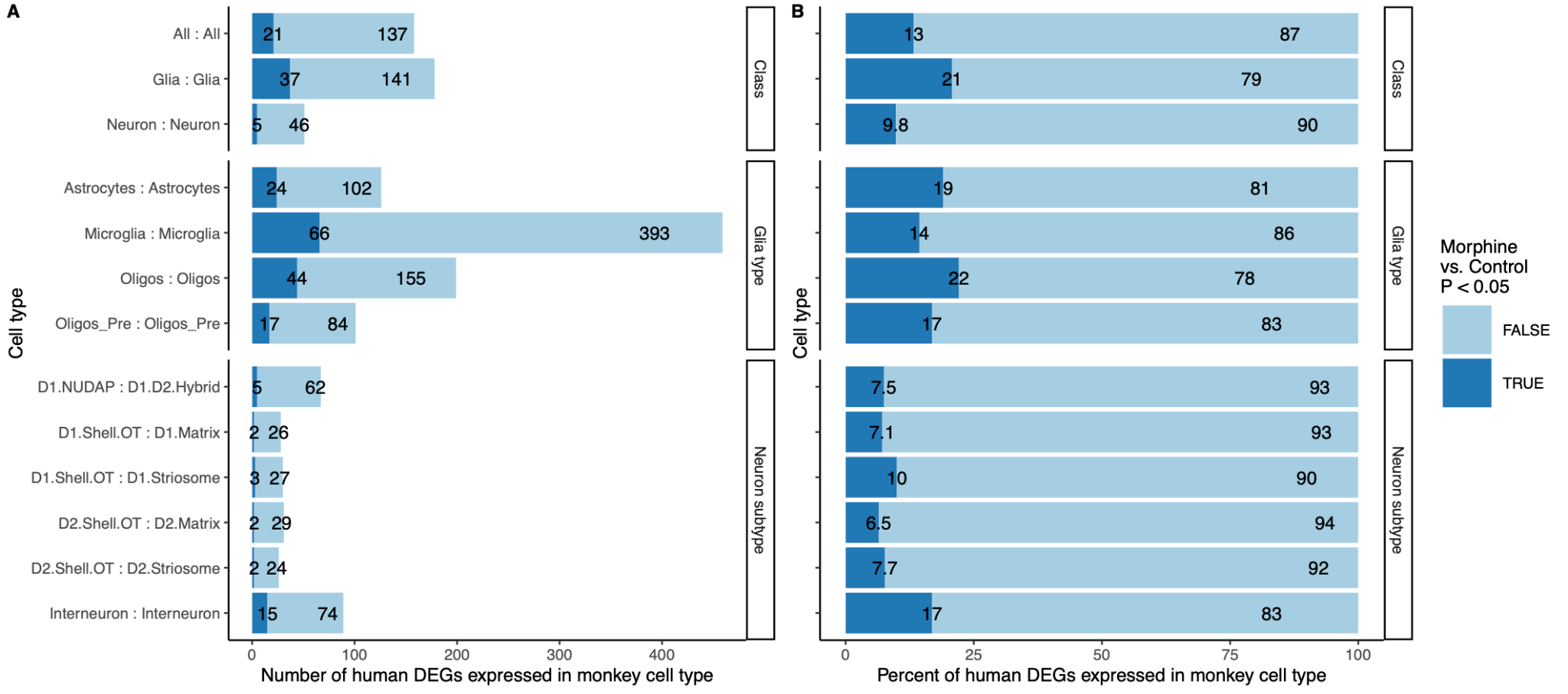

###### Figure S9. Replication of human opioid use disorder differential expression in rhesus macaque striatal cell types

Barplots in the number (A) or percent (B) of human differentially expressed genes (DEGs) in opioid use disorder within dorsal striatal cell types that are also differentially expressed at replication P < 0.05 in the corresponding rhesus macaque ventral striatal cell types in chronic morphine exposure.

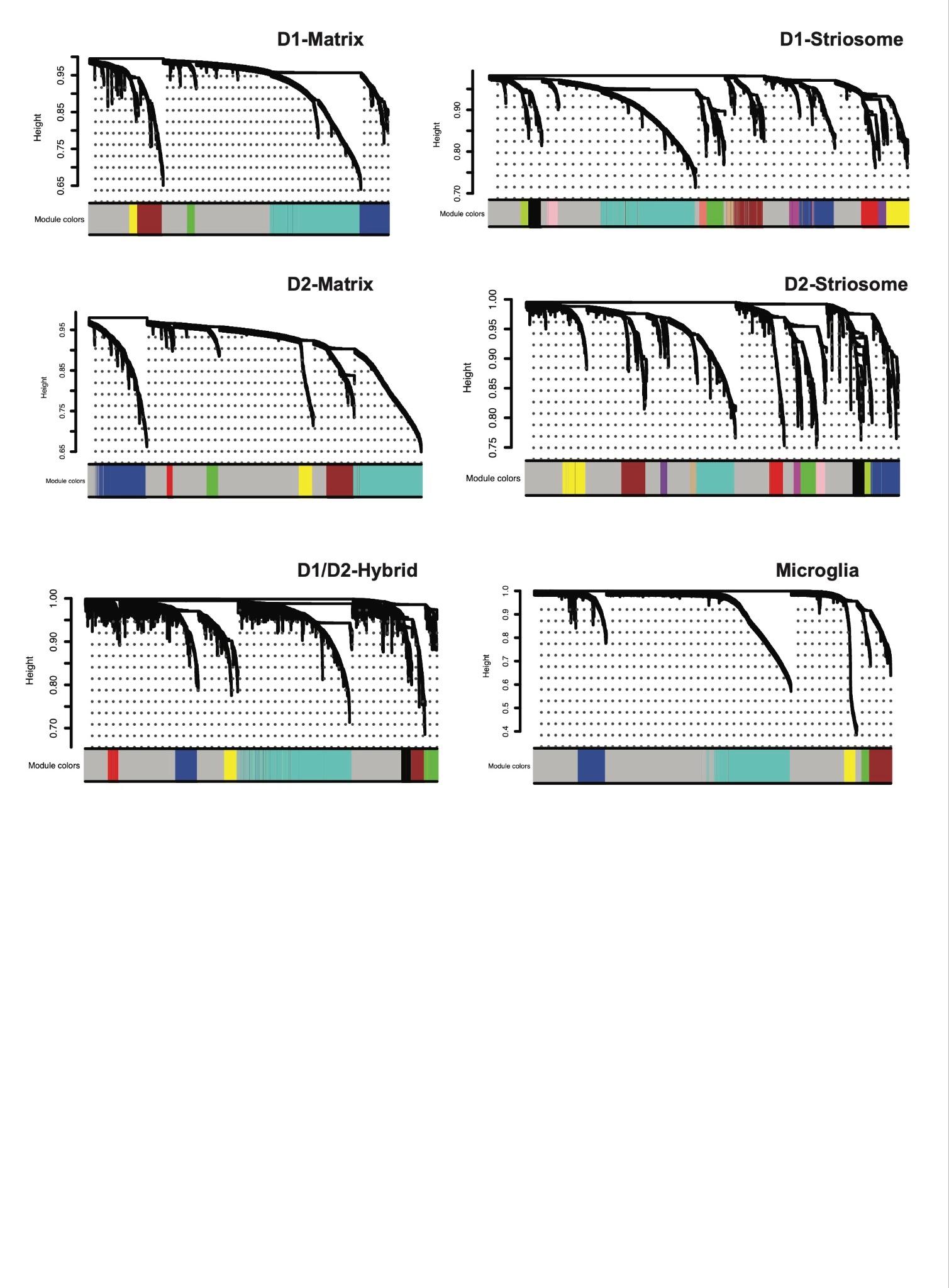

###### Figure S10. Dendrograms of gene co-expression networks among modules in medium spiny neuron cell types and microglia.

Dendrograms from WGCNA showing the average linkage hierarchical clustering of genes per cell type.

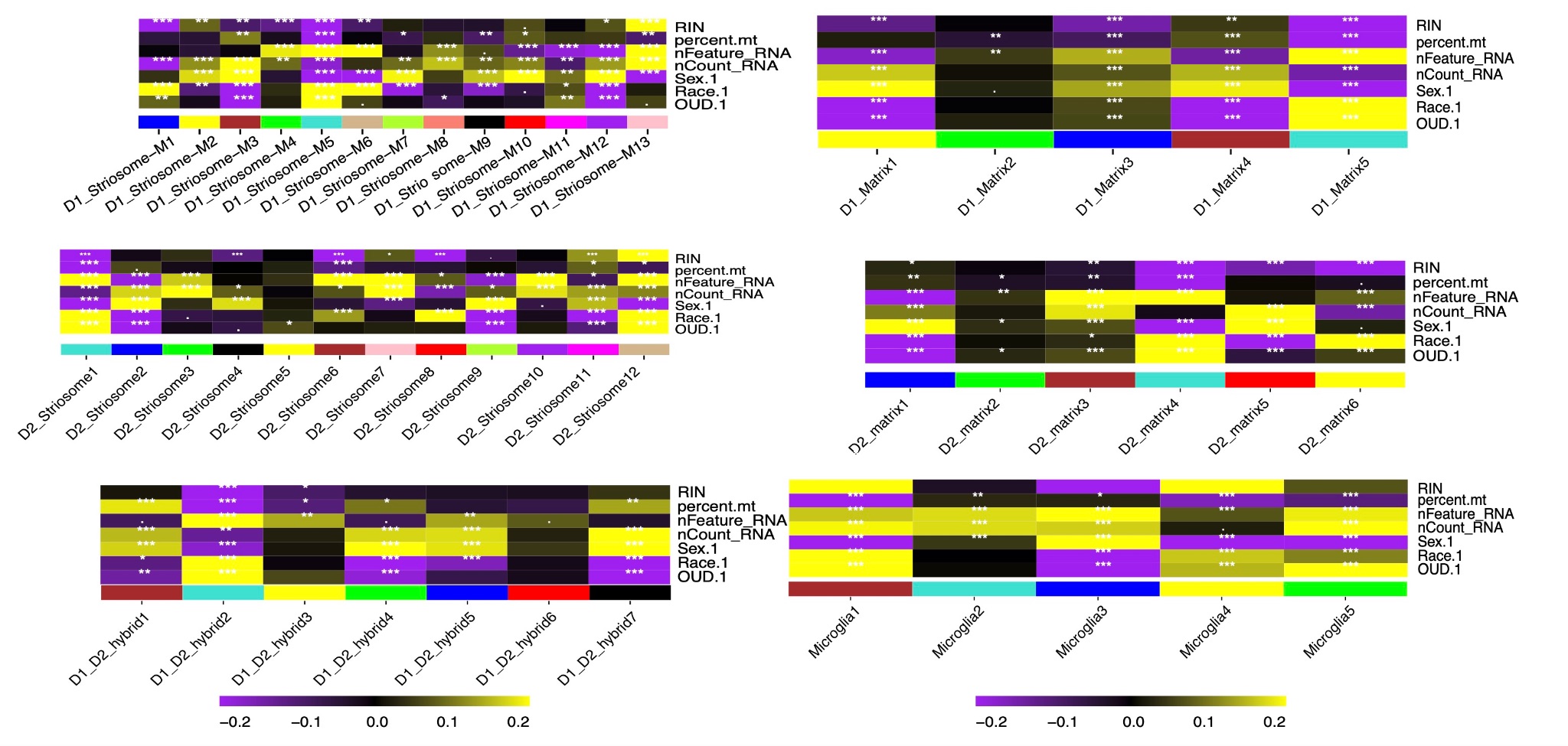

###### Figure S11. Module-trait relationships for striatal cell types.

Correlation between modules (except grey module) and each of 6 traits (RIN, percent_mt, nFeature_RNA, nCount_RNA, Sex, Race and OUD). Correlation coefficient value was suggested by color (purple: negative correlation, yellow: positive correlation). ***: FDR < 0.001, **: FDR < 0.01, *: FDR < 0.05.

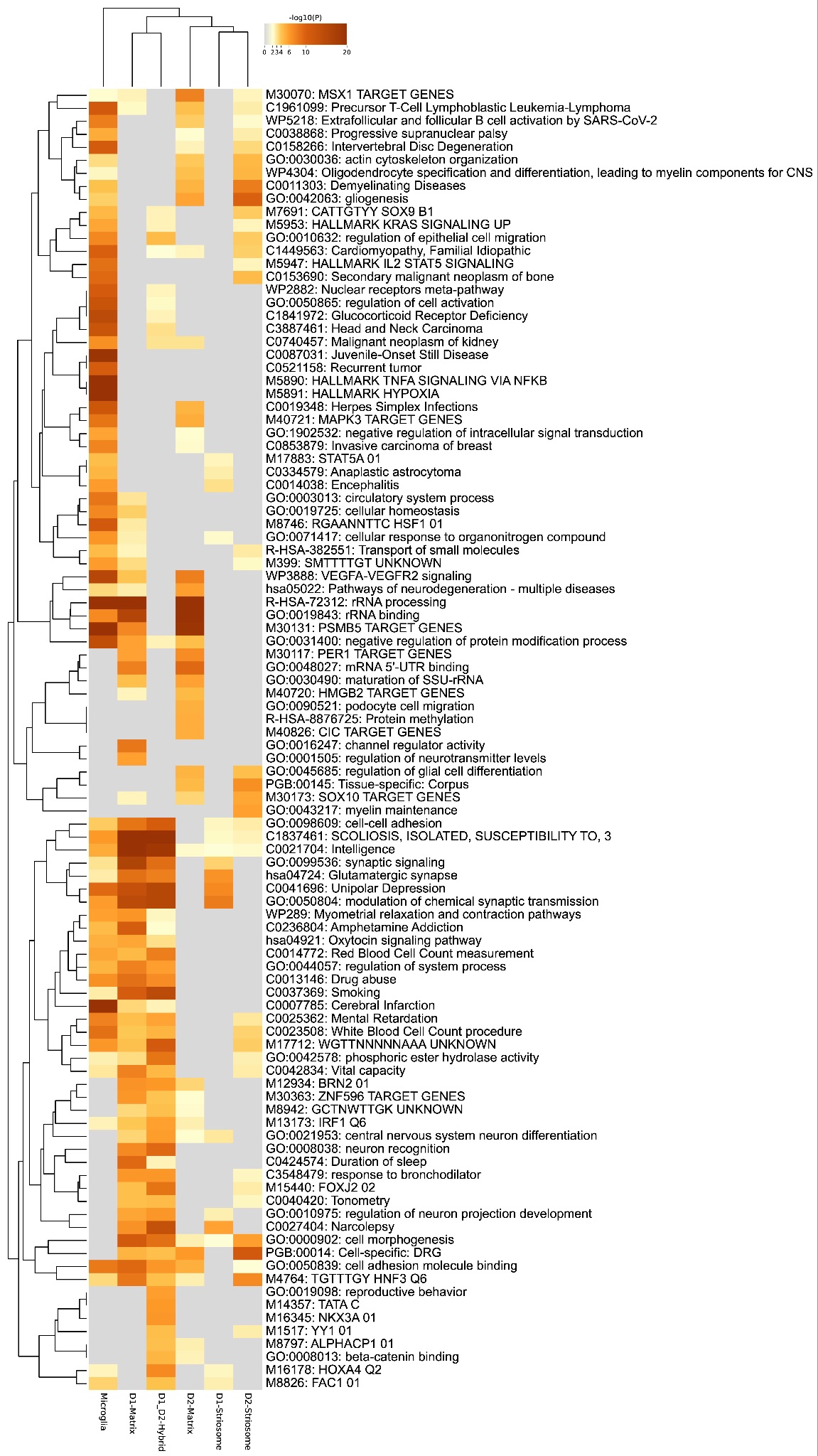

###### Figure S12. Top 100 enriched pathways among genes within significant OUD-associated modules in medium spiny neuron subpopulations and microglia.

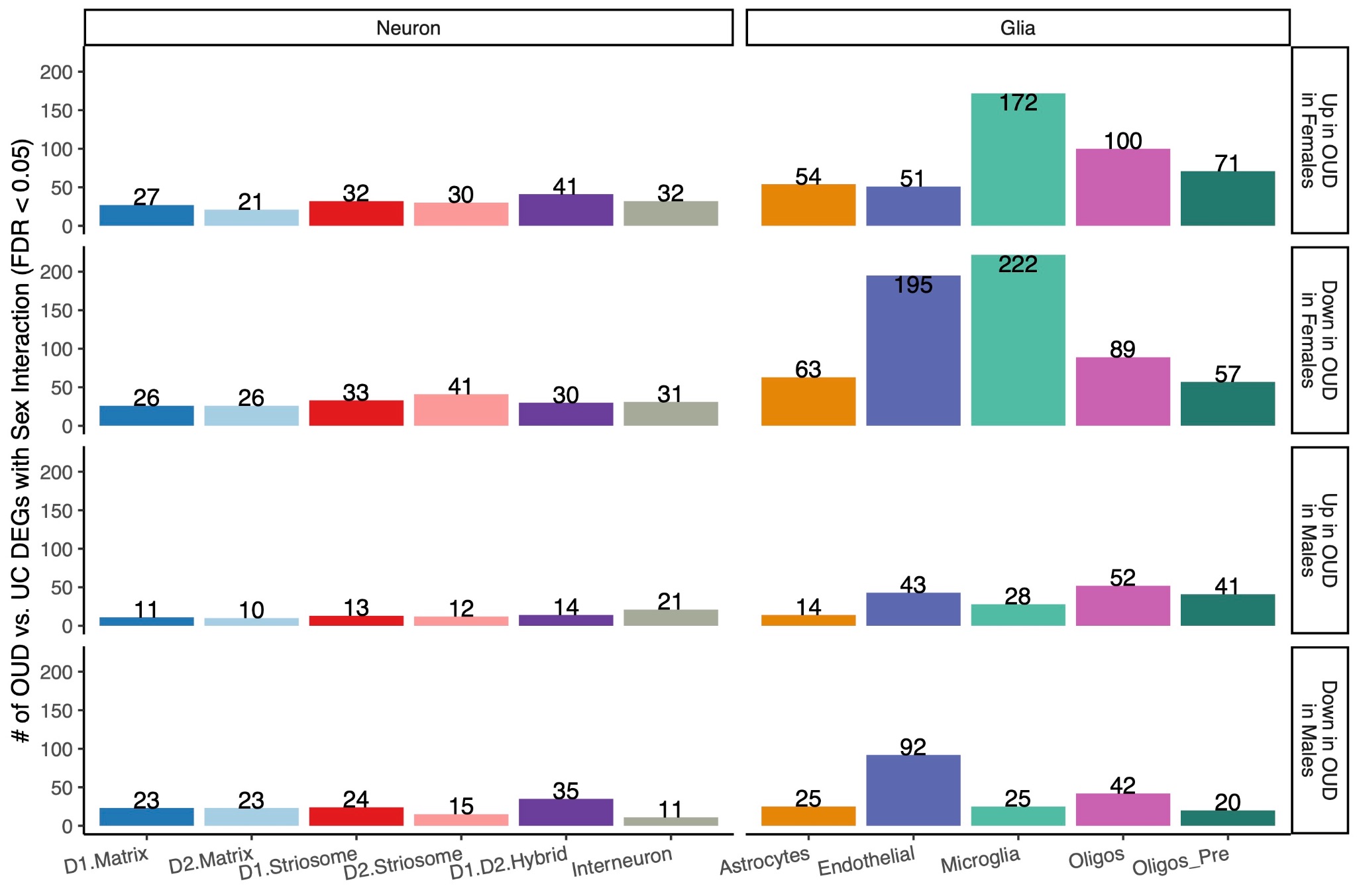

Figure S13. The effect of biological sex and OUD diagnosis in striatal cell types.

Barplot of striatal cell types with significant interaction effects (FDR < 0.05) stratified by cell type and main effects. The genes are grouped into four groups by the direction of the interaction fold-change and whether the OUD effect is larger in females or males.

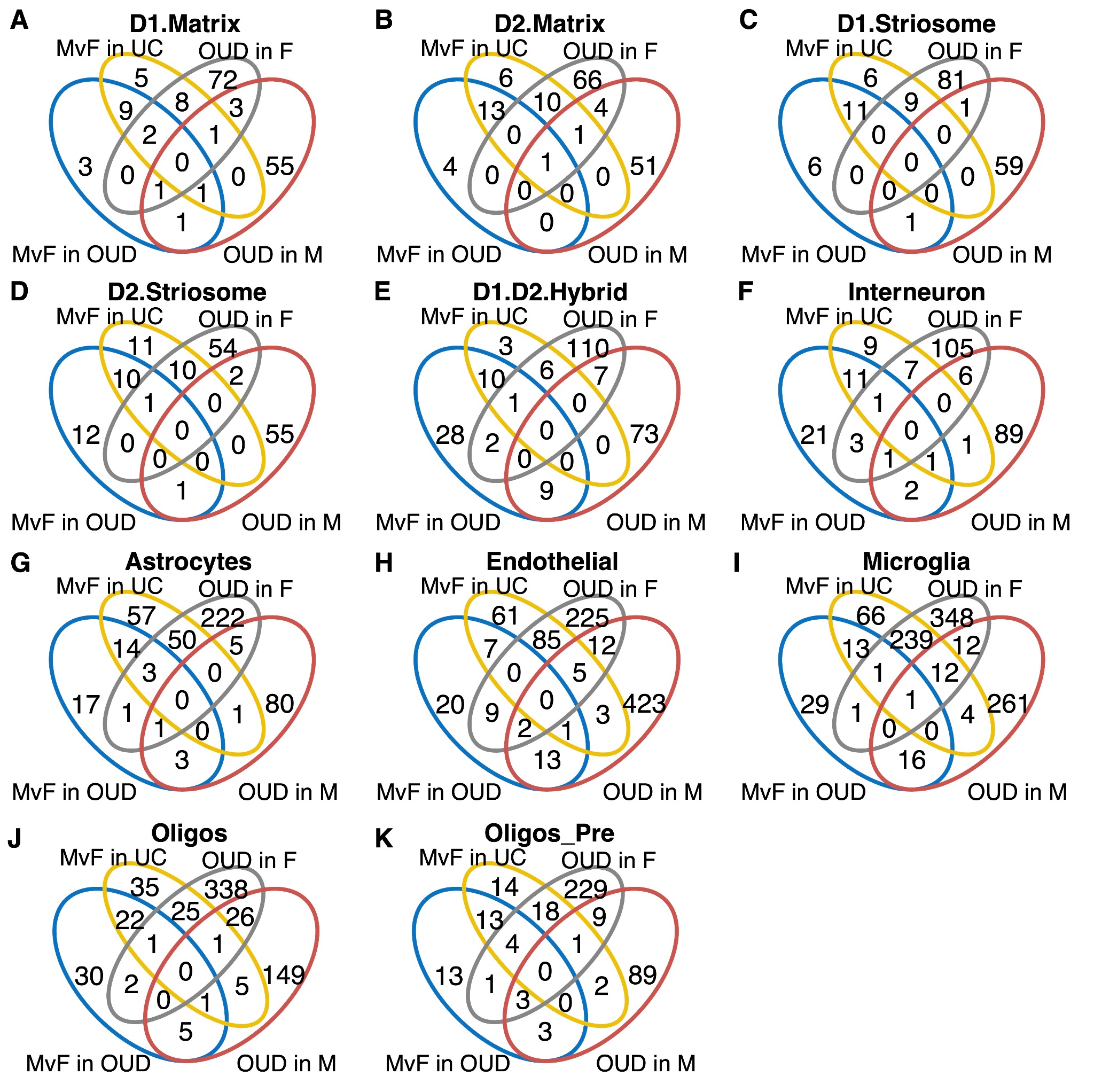

Figure S14. Limited overlap of sex-specific changes in OUD compared to unaffected individuals with differences between sexes in OUD or unaffected individuals.

Four-way venn diagram showing the number of overlapping differentially expressed genes in the following groups (FDR < 0.05) within striatal neuronal cell types (A-F) and glial cell types (G-K). MvF in UC (yellow oval): within the subset of unaffected individuals, the differentially expressed genes between females vs. males. MvF in OUD (blue oval): within the subset of OUD individuals, the differentially expressed genes between females vs. males (reference = male). OUD in F (gray oval): within the subset of female individuals, the differentially expressed genes between OUD vs. unaffected individuals. OUD in M (red oval): within the subset of male individuals, the differentially expressed genes between OUD vs. unaffected individuals.

##### Data and Code Availability

Single nuclei RNA-seq data processing and downstream analyses of the human dorsal striatum in this paper are collected in the github repository [https://github.com/pfenninglab/Logan_Striatum_snRNA-seq](https://github.com/pfenninglab/Logan_BU_Striatum_snRNA-seq). Single nuclei RNA-seq processing and downstream analyses of the rhesus macaque nucleus accumbens in this paper are collected in the github repository <https://github.com/pfenninglab/McLean_chronic_opioid_monkey_snRNA-seq>.The raw sequencing reads and annotated Seurat objects for both human and rhesus macaque studies are uploaded to GEO under SuperSeries accession number (XXXXXXX) accessible during review with the token qpyxymamptcbbij. A browsable webportal of the human dorsal striatum single nuclei transcriptomes are on the CZ CellxGene Discovery webportal at <https://cellxgene.cziscience.com/collections/cec4ef8e-1e70-49a2-ae43-1e6bf1fd5978>.

##### Inclusion & Diversity

One or more of the authors of this paper self-identifies as a member of the LGBTQ+ community.

**Acknowledgements**

Thank you to the staff and technicians who work diligently as part of the Brain Tissue Donation Program at the University of Pittsburgh. Postmortem human brain tissue was provided by the University of Pittsburgh Brain Tissue Donation Program and the National Institutes of Health NeuroBioBank at the University of Pittsburgh.

**Funding**

The research reported in this article was supported by: Research was supported by NIH HEAL Initiative under National Heart, Lung, and Blood Institute (R01HL150432, RWL); and National Institute on Drug Abuse (R01DA051390, MLS and RWL; DP1DA046585, ARP; F30DA053020, BNP; R01DA047130, SJK).

**Author Contributions**

RWL conceptualized and managed the project. RWL and ARP provided supervision for data analyses and interpretation. BNP, ARP, RWL, SJK, and MLS obtained funding for the project. BNP, MHR, JFG, DAL, ZF, MLW, ARP, and RWL designed and implemented the project. MHR, MKF, AET, SJR, and YA conducted human tissue homogenization, single nuclei extraction, and single nuclei library preparations. RJF, SJK, JB, SNH, KMM, and KJR conceptualized and managed the rhesus macaque project, including tissue homogenization, single nuclei extraction, and single nuclei library preparations. BNP and CF completed the computational analyses and BNP, MHR, QS, and RWL provided software, visualization and validation. BNP, XX, QS, GCT, ARP, and RWL designed and interpreted analyses and outcomes. RWL, JG, DAL, and ARP provided the tissue resources and computational infrastructure for analyses. BNP, MHR, MKF, ARP, and RWL wrote the original manuscript with figures. All authors participated in reviewing and editing the manuscript for publication.

### References (152-181 unique to Supplement)

1.	National Institute on Drug Abuse. Drug Overdose Death Rates. *National Institute on Drug Abuse* <https://nida.nih.gov/research-topics/trends-statistics/overdose-death-rates> (2023).

2.	Puig, S. *et al.* Uncovering circadian rhythm disruptions of synaptic proteome signaling in prefrontal cortex and nucleus accumbens associated with opioid use disorder. *bioRxiv* (2023) doi:[10.1101/2023.04.07.536056](http://dx.doi.org/10.1101/2023.04.07.536056).

20.	Madabhushi, R., Pan, L. & Tsai, L.-H. DNA damage and its links to neurodegeneration. *Neuron* **83**, 266–282 (2014).

21.	Shanbhag, N. M. *et al.* Early neuronal accumulation of DNA double strand breaks in Alzheimer’s disease. *Acta Neuropathologica Communications* vol. 7 Preprint at https://doi.org/[10.1186/s40478-019-0723-5](http://dx.doi.org/10.1186/s40478-019-0723-5) (2019).

43.	Gayden, J. *et al.* Three-dimensional characterization of medium spiny neuron heterogeneity in the adult mouse striatum. *bioRxiv* (2023) doi:[10.1101/2023.05.04.539488](http://dx.doi.org/10.1101/2023.05.04.539488).

55.	Mateusz, S. L. *et al.* β-catenin signaling via astrocyte-encoded TCF7L2 regulates neuronal excitability and social behavior. *bioRxiv* 2020.11.28.402099 (2020) doi:[10.1101/2020.11.28.402099](http://dx.doi.org/10.1101/2020.11.28.402099).

150.	Nylander, I. *et al.* Evidence for a Link Between Fkbp5/FKBP5, Early Life Social Relations and Alcohol Drinking in Young Adult Rats and Humans. *Mol. Neurobiol.* **54**, 6225–6234 (2017).

151.	Glausier, J. R., Kelly, M. A., Salem, S., Chen, K. & Lewis, D. A. Proxy measures of premortem cognitive aptitude in postmortem subjects with schizophrenia. *Psychol. Med.* **50**, 507–514 (2020).

152.	Mai, J. K., Majtanik, M. & Paxinos, G. *Atlas of the Human Brain*. (Academic Press, 2015).

153.	National Research Council, Division on Earth and Life Studies, Institute for Laboratory Animal Research & Committee for the Update of the Guide for the Care and Use of Laboratory Animals. *Guide for the Care and Use of Laboratory Animals: Eighth Edition*. (National Academies Press, 2011).

154.	Holtzman, S. G. & Villarreal, J. E. Operant behavior in the morphine-dependent rhesus monkey. *J. Pharmacol. Exp. Ther.* **184**, 528–541 (1973).

155.	Borgatti, D., Bergman, J. & Kohut, S. Effects of lorcaserin on opioid‐induced observable behavior in rhesus monkeys. *FASEB J.* **34**, 1–1 (2020).

156.	Renthal, W. *et al.* Characterization of human mosaic Rett syndrome brain tissue by single-nucleus RNA sequencing. *Nat. Neurosci.* **21**, 1670–1679 (2018).

157.	Klein, A. M. *et al.* Droplet barcoding for single-cell transcriptomics applied to embryonic stem cells. *Cell* **161**, 1187–1201 (2015).

158.	Kaminow, B., Yunusov, D. & Dobin, A. STARsolo: accurate, fast and versatile mapping/quantification of single-cell and single-nucleus RNA-seq data. *bioRxiv* 2021.05.05.442755 (2021) doi:[10.1101/2021.05.05.442755](http://dx.doi.org/10.1101/2021.05.05.442755).

159.	Shumate, A. & Salzberg, S. L. Liftoff: accurate mapping of gene annotations. *Bioinformatics* **37**, 1639–1643 (2021).

160.	Phan, B. & Pfenning, A. Alternate gene annotations for rat, macaque, and marmoset for single cell RNA and ATAC analyses. *https://kilthub.cmu.edu/articles/dataset/Alternate_gene_annotations_for_rat_macaque_and_marmoset_for_single_cell_RNA_and_ATAC_analyses/21176401* <https://doi.org/10.1184/R1/21176401.v1> (2022) doi:[10.1184/R1/21176401.v1](http://dx.doi.org/10.1184/R1/21176401.v1).

161.	Young, M. D. & Behjati, S. SoupX removes ambient RNA contamination from droplet-based single-cell RNA sequencing data. *Gigascience* **9**, (2020).

162.	Muskovic, W. & Powell, J. E. DropletQC: improved identification of empty droplets and damaged cells in single-cell RNA-seq data. *Genome Biol.* **22**, 1–9 (2021).

163.	Bais, A. S. & Kostka, D. scds: computational annotation of doublets in single-cell RNA sequencing data. *Bioinformatics* **36**, 1150–1158 (2019).

164.	Hippen, A. A. *et al.* miQC: An adaptive probabilistic framework for quality control of single-cell RNA-sequencing data. *PLoS Comput. Biol.* **17**, e1009290 (2021).

165.	Hao, Y. *et al.* Integrated analysis of multimodal single-cell data. *Cell* **184**, 3573–3587.e29 (2021).

166.	Hafemeister, C. & Satija, R. Normalization and variance stabilization of single-cell RNA-seq data using regularized negative binomial regression. *Genome Biol.* **20**, 296 (2019).

167.	Ahlmann-Eltze, C. & Huber, W. glmGamPoi: fitting Gamma-Poisson generalized linear models on single cell count data. *Bioinformatics* **36**, 5701–5702 (2020).

168.	Crowell, H. L. *et al.* muscat detects subpopulation-specific state transitions from multi-sample multi-condition single-cell transcriptomics data. *Nat. Commun.* **11**, 1–12 (2020).

169.	Soneson, C. & Robinson, M. D. Bias, robustness and scalability in single-cell differential expression analysis. *Nat. Methods* (2018).

170.	Squair, J. W. *et al.* Confronting false discoveries in single-cell differential expression. *Nat. Commun.* **12**, 5692 (2021).

171.	Robinson, M. D., McCarthy, D. J. & Smyth, G. K. edgeR: a Bioconductor package for differential expression analysis of digital gene expression data. *Bioinformatics* **26**, 139–140 (2009).

172.	Subramanian, A. *et al.* Gene set enrichment analysis: a knowledge-based approach for interpreting genome-wide expression profiles. *Proc. Natl. Acad. Sci. U. S. A.* **102**, 15545–15550 (2005).

173.	Korotkevich, G. *et al.* Fast gene set enrichment analysis. *bioRxiv* 060012 (2021) doi:[10.1101/060012](http://dx.doi.org/10.1101/060012).

174.	Koopmans, F. *et al.* SynGO: An Evidence-Based, Expert-Curated Knowledge Base for the Synapse. *Neuron* **103**, 217–234.e4 (2019).

175.	Kowalczyk, A., Chikina, M. & Clark, N. Complementary evolution of coding and noncoding sequence underlies mammalian hairlessness. *Elife* **11**, (2022).

176.	Kowalczyk, A., Partha, R., Clark, N. L. & Chikina, M. Pan-mammalian analysis of molecular constraints underlying extended lifespan. *Elife* **9**, (2020).

177.	Csárdi, G. & Nepusz, T. The igraph software package for complex network research. (2006).

178.	Van de Sande, B. *et al.* A scalable SCENIC workflow for single-cell gene regulatory network analysis. *Nat. Protoc.* **15**, 2247–2276 (2020).

179.	Moerman, T. *et al.* GRNBoost2 and Arboreto: efficient and scalable inference of gene regulatory networks. *Bioinformatics* **35**, 2159–2161 (2018).

180.	Boca, S. M. & Leek, J. T. A direct approach to estimating false discovery rates conditional on covariates. *PeerJ* **6**, e6035 (2018).

181.	Korthauer, K. *et al.* A practical guide to methods controlling false discoveries in computational biology. *Genome Biol.* **20**, 1–21 (2019).
